# The Notch signaling pathway is a master regulator of CD8^+^ T cell exhaustion and differentiation during chronic infection

**DOI:** 10.64898/2026.09.01.748651

**Authors:** Dave Maurice De Sousa, Salix Boulet, Frédéric Duval, Eric Perkey, Jean-François Daudelin, Laure Le Corre, Marie-Ève Lebel, Alain Lamarre, Freddy Radtke, Burkhard Ludewig, Chris Siebel, Ivan Maillard, Nathalie Labrecque

## Abstract

During chronic infection, the persistence of antigen and inflammation leads to the differentiation of CD8^+^ T cells into an exhausted state characterised by expression of inhibitory receptors (IRs) and the progressive loss of T cell functions. Among the different subsets of exhausted CD8^+^ T (Tex) cells, Tex progenitors expressing SLAMF6 and the transcription factor TCF-1 (TCF-1+) give rise to Tex effector-like cells expressing CX3CR1 and Tex terminal cells expressing CD101. PD-1/PD-L1 blockade acts on TCF-1+ Tex progenitor cells and promotes their differentiation into Tex effector-like cells. The molecular events controlling CD8^+^ Tex cell differentiation are still poorly defined. As Notch signaling may be sustained during chronic infection by persistent TCR stimulation and inflammation, we tested whether Notch signaling influences CD8^+^ T cell exhaustion. Using mice lacking (N1N2^Δ/Δ^) or not (N1N2^fl/fl^) Notch1/2 expression only in mature CD8^+^ T cells, we showed that the absence of Notch signal causes severe CD8+ T cell exhaustion during chronic LCMV infection. N1N2^Δ/Δ^ Tex cells express higher levels of IRs and are less functional when compared to their wild-typee counterpart. In the absence of N1N2 receptors, Tex progenitor and Tex terminal cells accumulate and Tex cells cannot be reinvigorated by PD-1/PD-L1 blockade. We further demonstrated that Notch signaling is essential to promote the differentiation of Tex progenitors into Tex effector-like cells. Moreover, Notch signals, provided by stromal cells expressing the ligands Delta-like 1 and 4, are necessary during all stages of the infection to prevent severe exhaustion. Single-nucleus RNA and ATAC multiome profiling identifies Notch signaling as a critical role on effector transcriptional programming in exhausted CD8 T cells. Loss of Notch signaling impairs transcriptional program associated with migration and perception of CD4^+^ T cell help. Together, these alterations drive the differentiation of Tex progenitor cells toward a terminally exhausted Tex fate.

## INTRODUCTION

Following an acute infection, CD8^+^ T cells proliferate and differentiate into short-lived effector cells (SLEC) or memory precursor effectors cells (MPEC) to control the infection. During the contraction phase, 90% of the cells, especially the SLECs, will die by apoptosis while MPECs will differentiate into functional memory T cells to protect the body against future re-infection^1–3^ . If the infection becomes chronic or in cancer, antigen persistence and sustained inflammation divert the differentiation process. In these conditions, CD8^+^ T cells will progressively acquire expression of inhibitory receptors (IRs), suffer from a gradual loss of function and effector CD8^+^ T cells will not differentiate into functional memory T lymphocytes but into exhausted CD8^+^ T cells (Tex)^4,5^ .

The use of monoclonal antibodies (mAbs) blocking IRs, such anti-PD-1/PD-L1 mAbs, has been shown to reinvigorate CD8^+^ Tex cells, restoring their capacity to eliminate infected and cancer cells^6^.Not all CD8^+^ Tex cells are reinvigorated following anti-PD-1/PD-L1 treatment, as different subsets of CD8+ Tex cells are generated following chronic infection with lymphocytic choriomeningitis virus (LCMV) clone 13 or in cancer. These subsets have different functions and sensitivity to anti-PD-1/PD-L1 therapy^7–10^. CD8+ Tex stem-like cells or Tex progenitor cells are characterized by elevated expression of SLAMF6 and TCF-1 and low expression of TIM-3 and CX3CR1^7,9,11,12^. They are responsible for maintaining the immune response throughout chronic infection and are the main subset responding to checkpoint blockade^11,12^. Tex stem-like cells can be further divided into two populations, SLAMF6^hi^ (TCF-1+) CD8^+^ Tex cells expressing CD69 (Tex prog1) located in the white pulp of the spleen while SLAMF6^hi^ CD69^lo^ CD8^+^ Tex cells (Tex prog2) found in the red pulp and blood^8,10^. They can produce cytokines such as IFN-γ but are poorly cytotoxic. Tex-prog cells will differentiate into Tex effector-like (Tex eff-like) cells characterized by the loss of SLAMF6, CD69 and TCF-1 expression and the gain of TIM-3, CX3CR1 and T-bet expression^7–9^. These cells are known to have better effector functions and efficiently produce IFN-γ and granzymeB (GzmB) to control viral load^7–9^. Also, optimal generation of these cells requires the help of CD4^+^ T cells ^7,13^. Tex-eff cells are mainly found in the circulation and infected tertiary tissues^8^. Tex-eff cells can further differentiate into Tex terminal (Tex-term) cells, which are mainly found in infected organs including secondary lymphoid organs (SLO)^8^, express more IRs such as PD-1 and TIM-3, acquire expression of CD101 ^9^, can re-express CD69^8^, lose expression of the transcription factor T-bet and are poorly functional^7–9^. The linear differentiation of Tex-prog into Tex-eff giving rise to Tex-term cells has been challenged by others that have shown poor differentiation of Tex-eff into Tex-term following adoptive transfer^14^ . Understanding which factors influence the differentiation, maintenance and survival of the different Tex populations is needed to optimize the efficacy of cell-based therapeutic approaches to control chronic infection and cancer.

The Notch signaling pathway controls many differentiation choices in the immune system^15^. In CD8^+^ T cells, we have shown that Notch signaling plays a key role in the differentiation of CD8^+^ T cells into SLECs following acute infection^16–18^. The Notch pathway is activated when one of the receptors (Notch1 to 4) interacts with its ligand (Jagged1, 2; Delta-like (DLL) 1, 3, 4) leading to the cleavage of the intracellular domain of Notch that will migrate to the nucleus to induce the transcription of target genes specific to the Notch pathway, such as Hes1 and Dtx1, but also genes specific to each cell type^19,20^. In mature CD8^+^ T cells, expression of Notch1 (N1) and Notch2 (N2) receptors is induced following stimulation of T cells via antigen and inflammatory cytokines^17,21^. In addition, ligands that activate the Notch pathway are expressed by antigen presenting cells and SLO stromal cells^22^. We therefore hypothesized that during a chronic infection, the persistence of the antigen and sustained inflammatory signals will lead to sustained activation of the Notch signaling pathway impacting CD8^+^ T cell fate.

In this paper, we show that early and continuous Notch signal, via DLL1/4 expressing stromal cells of SLOs, prevents severe exhaustion of CD8^+^ T cells. Furthermore, we demonstrate that Notch is essential for the reinvigoration of CD8^+^ T cells following anti-PD-L1 therapy and for the differentiation of Tex-prog cells into Tex-eff cells. Loss of N1N2 receptors specifically on CD8^+^ T cells causes severe exhaustion and the accumulation of CD8^+^ Tex-term cells, which impairs the control of viral load following chronic infection. Overall, our results show that the Notch signaling pathway is a central regulator of CD8^+^ T cell exhaustion.

## RESULTS

### The Notch signaling pathway prevents severe CD8^+^ T cell exhaustion

The Notch signaling pathway plays important role during acute CD8^+^ T cell response^16,17^, whether it also influences CD8^+^ T cell response during chronic infection is a likely possibility as TCR stimulation and inflammatory signals are known inducers of Notch receptor expression in T cells and of Notch ligands on dendritic cells^17,18,21^. To ask whether Notch signaling regulates the exhaustion state of CD8^+^ T cells during chronic infection, we used mice in which Notch1 an Notch2 expression is only abrogated in mature peripheral CD8^+^ T cells^16^. *Notch1* and *Notch2* floxed mice (N1N2^fl/fl^) were crossed with E8I-cre+ mice to generate mice in which Notch1 and Notch2 are inactivated only in mature CD8^+^ T cells (N1N2^Δ/Δ^). Following infection of these mice with LCMV clone 13, we observed an increase in the number of gp33-specific CD8^+^ T cells in N1N2^Δ/Δ^ mice compared to their littermate controls (N1N2^fl/fl^) (Fig. 1A). A better response of Notch-deficient CD8^+^ T cells was also observed at day 8 and day 21 post-infection (Fig. S1A).

**Figure 1.**
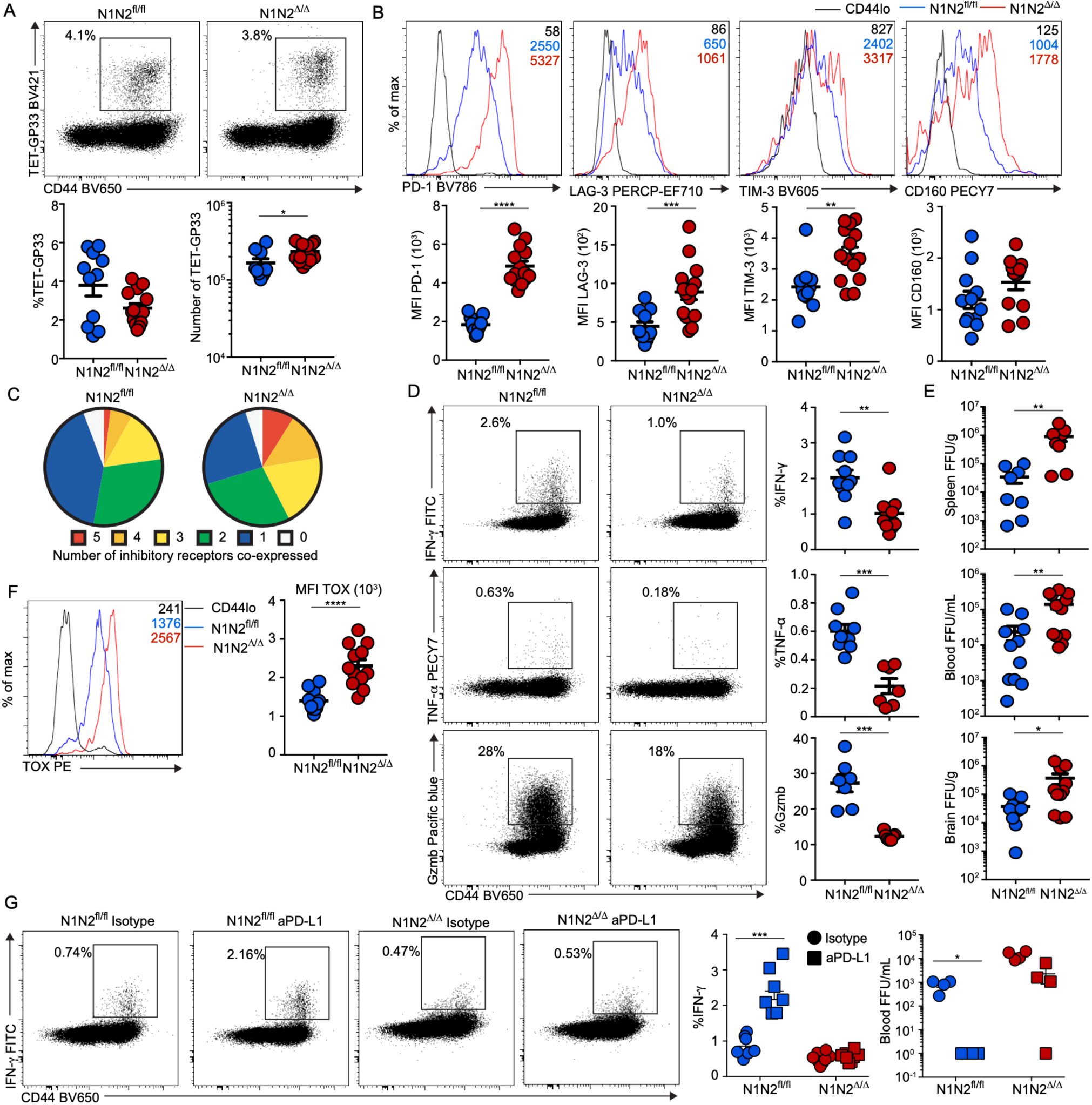
Notch signaling limits severe exhaustion during chronic infection. (**A-C**) Mice were infected with LCMV clone 13 and the CD8^+^ T cell response in the spleen was characterized at d30 post-infection by flow cytometry via tetramer staining. (**A**) Percentage and numbers of Tet-gp33 in CD8^+^ T cells from N1N2^fl/fl^ and N1N2^Δ/Δ^ mice. (**B**) Expression of inhibitory receptors on LCMV clone 13-specific CD8^+^ T cells from N1N2^fl/fl^ and N1N2^Δ/Δ^ mice (**C**) Co-expression of inhibitory receptors (PD-1, LAG-3, TIM-3, CD160 and 2B4) on gp33-specific CD8^+^ T cells from N1N2^fl/fl^ and N1N2^Δ/Δ^ mice on a per cell basis. (**D**) At d30 post-infection, cytokine production by LCMV-specific CD8^+^ T cells from N1N2^fl/fl^ and N1N2^Δ/Δ^ mice was measured ex vivo after a 5 hours restimulation with the gp33 peptide. **(E)** Viral loads in spleen, blood, and brain of N1N2^fl/fl^ and N1N2^Δ/Δ^ at d30 post-infection were determined using focus-forming assay on MC57G cells. **(F)** Expression of transcription factor TOX on Tet-gp33^+^ Notch-sufficient and -deficient CD8^+^ T cells. **(G)** N1N2^fl/fl^ and N1N2^Δ/Δ^ mice were infected with LCMV clone 13 and treated with 0.2 mg of anti-PD-L1 or isotype control every 3 days between d23 and d35 post-infection. At d37, IFN-γ production by gp33-restimulated CD8^+^ T cells was characterized and blood viral load determined. Data are from three (A-F) or two (G) independent experiments. Unpaired Student’s *t* test, with a Welch’s correction when applied, was used for two-group comparison and Kruskal–Wallis ANOVA with Dunn’s multiple comparison for multiple group comparison. \**P*<0.05, \*\**P*<0.01, \*\*\**P*<0.001, \*\*\*\**P*<0.0001.

In absence of Notch signaling, gp33-specific CD8^+^ T cells express higher levels of IRs such as PD-1, TIM-3, CD160 and LAG-3 at day 30 post-infection (Fig. 1B). PD-1 expression is also higher on N1N2^Δ/Δ^ gp33-specific CD8^+^ T cells than their wild-type counterparts at day 8 and day 21 post-infection (Fig. S1B). On a per cell basis, N1N2^Δ/Δ^ gp33-specific CD8^+^ T cells express more IRs compared to N1N2^fl/fl^ CD8^+^ T cells (Fig. 1C). Ex-vivo restimulation of CD8^+^ T cells using gp33 peptide showed that the proportion of CD8^+^ T cells producing IFN-γ, TNF-α and GzmB is significantly reduced in N1N2^Δ/Δ^ CD8^+^ T cells compared to their wild-type counterpart (Fig. 1D). This loss of functionality correlates with higher viral load in the blood, spleen and brain at day 30 post-infection (Fig. 1E). The increase in viral load is not observed at d8 but is already higher at d21 post-infection in the blood (Fig. S1C). The master transcriptional regulator of CD8^+^ T cell exhaustion, TOX^23–26^, was also expressed at higher level in N1N2^Δ/Δ^ than in N1N2^fl/fl^ gp33-specific CD8^+^ T cells at day 8, 21 and 30 post-infection (Fig. 1F and Fig. S1D). To validate that the severe exhaustion of CD8^+^ T cells induced by lack of Notch signals was not a consequence of increased viral load, we adoptively transferred N1N2^fl/fl^ or N1N2^Δ/Δ^ P14 TCR transgenic CD8^+^ T cells into congenic B6.SJL recipients followed by infection with LCMV clone 13. At day 30, the severely exhausted phenotype was only observed in P14N1N2^Δ/Δ^ CD8^+^ T cells, even if the viral load was equal between recipient mice (Fig. S1E-G).

### Severely exhausted N1N2^Δ/Δ^ CD8^+^ T cells cannot be reinvigorated by anti-PD-L1 treatment

The severe CD8^+^ T cell exhaustion induced by Notch-deficiency led us to evaluate their response to anti-PD-L1 treatment. N1N2^fl/fl^ and N1N2^Δ/Δ^ mice were infected with LCMV clone 13 and treated with either isotype or anti-PD-L1 antibodies every 3 days between day 23 and day 35 post-infection. At day 37 post-infection, anti-PD-L1 treatment resulted in improved functions of N1N2^fl/fl^ gp33-specific CD8^+^ T cells as measured by increased IFN-γ production (Fig. 1G). However, this was not observed in the N1N2^Δ/Δ^ mice (Fig. 1G). Furthermore, viral load was only reduced in N1N2^fl/fl^ mice following anti-PD-L1 treatment (Fig. 1G). Therefore, in the absence of expression of the Notch1 and Notch2 receptors, severely exhausted CD8^+^ T cells cannot be reinvigorated by checkpoint blockade.

### Notch signaling acts as a master regulator of exhausted CD8^+^ T cell differentiation

Recent studies have described that exhausted CD8^+^ T cells are composed of different subsets that play different roles during exhaustion. First, using CD69 and SLAMF6 staining we observed increased proportion and numbers of the two progenitor subsets (Tex prog1 and Tex prog2) in absence of Notch signaling at day 30 post-infection (Fig. 2A). Furthermore, the proportion and the numbers of gp33-specific CD8^+^ T cells expressing TCF-1, an important transcription factor defining progenitor cells^11,12^, is higher in the absence of Notch signal. (Fig. 2B). On the other hand, the proportion and numbers of the effector-like subset (Tex Eff-like; CD69^lo^/SLAMF6^lo^) is severely decreased in exhausted N1N2^Δ/Δ^ CD8^+^ T cells compared to N1N2^fl/fl^ CD8^+^ T cells (Fig. 2A). In agreement with this observation, we observed a significant decrease in the proportion and numbers of CX3CR1^+^ Tex-eff cells in absence of Notch signaling (Fig. 2C)^7,9^. In addition, there was a significant accumulation in the proportion and numbers of terminally differentiated exhausted CD8^+^ T cells (Tex-term; SLAMF6^lo^/CD69^+^) (Fig. 2A) and CD101^+^ (another marker related to terminally exhausted cells) exhausted CD8^+^ T cells (Fig. 2C) in absence of Notch signaling in CD8^+^ T cells. It is important to note that these phenotypes are not dependent on the viral load as similar results were obtained in the P14 adoptive transfer model (Fig. S2). Furthermore, the differences between the proportion of Tex subsets are observed as soon as day 8 and day 21 post-infection (Fig. S3A-B). Altogether, these results suggest that Notch signaling is required for the differentiation of Tex-prog cells into Tex-eff cells but not for their differentiation into Tex-term cells.

**Figure 2.**
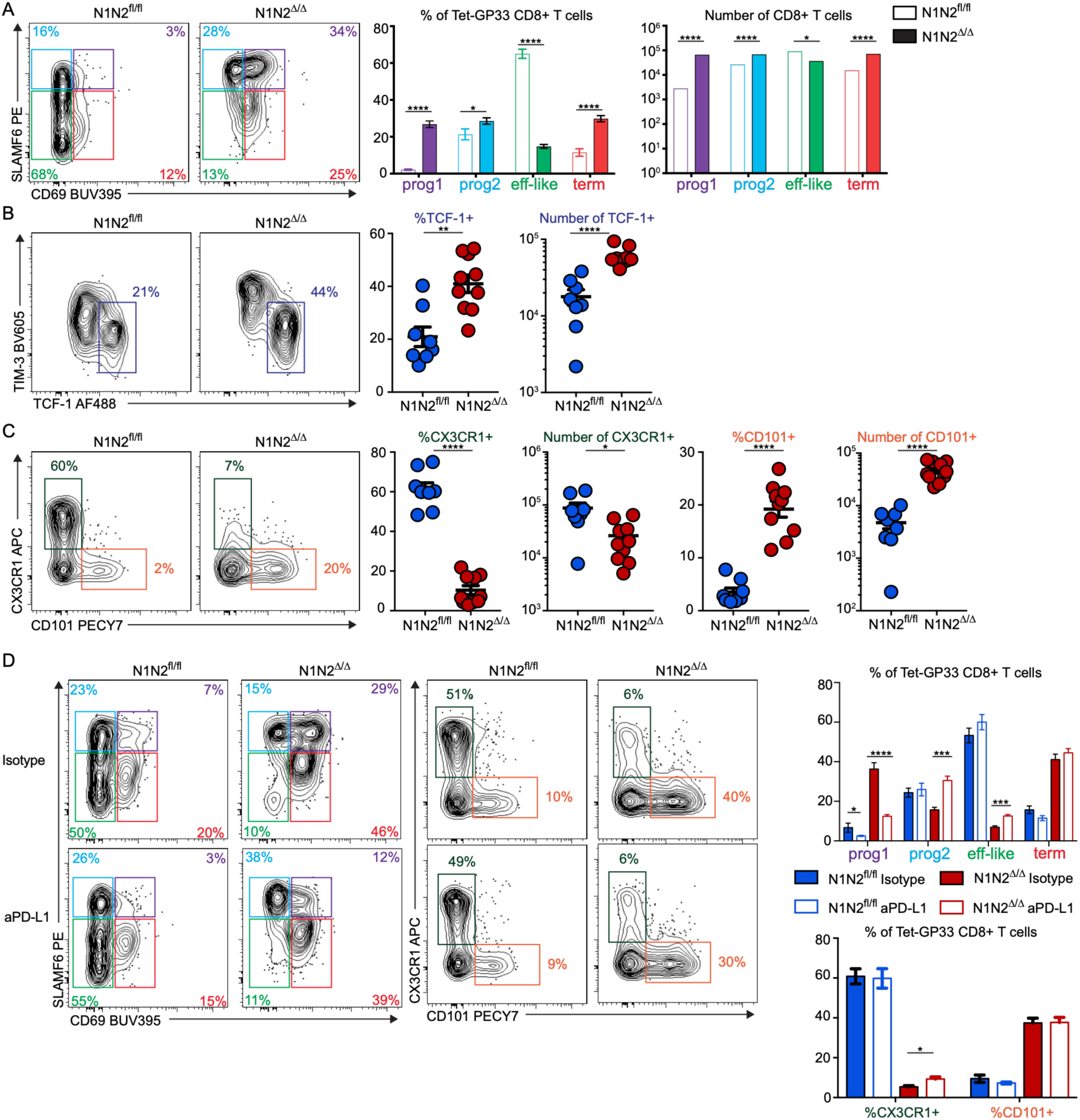
Notch signaling acts as a master regulator of exhausted CD8^+^ T cell differentiation. Mice were infected with LCMV clone 13 and the response in the spleen was characterized at d30 post-infection by flow cytometry via tetramer staining. The proportion and number of Tex subsets defined by CD69 and SLAMF6 (**A**), TCF-1 and Tim3 (**B**) and CD101 and CX3CR1 (**C**) was measured on gp33-specific CD8^+^ T cells by flow cytometry in N1N2^fl/fl^ and N1N2^Δ/Δ^ mice. (**D**) N1N2^fl/fl^ and N1N2^Δ/Δ^ mice were infected with LCMV clone 13 and treated with 0.2 mg of anti-PD-L1 or isotype control every 3 days between day 23 and day 35 post-infection. Proportion of Tex subsets, CX3CR1^+^ and CD101^+^ cells were determined on gp33-specific CD8+ T cells from N1N2^fl/fl^ and N1N2^Δ/Δ^ mice at day 37 post-infection. Data are from three (A-C) independent experiments. (D) is representative of 1 from 2 independent experiment. Unpaired Student’s *t* test, with a Welch’s correction when applied, was used for two-group comparison and Kruskal–Wallis ANOVA with Dunn’s multiple comparison for multiple group comparison. \**P*<0.05, \*\**P*<0.01, \*\*\**P*<0.001, \*\*\*\**P*<0.0001.

It is intriguing that Notch-deficient exhausted CD8^+^ T cells cannot be reinvigorated with anti-PD-L1 treatment even though they have a higher proportion of Tex prog cells in comparison to their wild-type counterparts^7–9^. To gain insight into this, we analysed the impact of anti-PD-L1 treatment on the distribution of the exhausted CD8^+^ T cell subsets in N1N2^Δ/Δ^ and N1N2^fl/fl^ mice. Following anti-PD-L1 treatment, N1N2^fl/fl^ and N1N2^Δ/Δ^ Tex prog1 cells responded to the treatment as seen by a decrease in their proportion while the proportion of Tex prog2 cells only increase in N1N2^Δ/Δ^ mice (Fig. 2D). Although the proportion of Tex eff-like cells increased in N1N2^Δ/Δ^ mice following anti-PD-L1 treatment, it does not reach the levels observed in N1N2^fl/fl^ mice (Fig. 2D). Although others have reported that anti-PD-L1 increases the generation of Tex-eff-like cells^27^, we did not observe such an increase in wild-type mice in the spleen. However, we did observe an increase in CX3CR1^+^ Tex eff-like cells in the liver of wild-type mice but in N1N2^Δ/Δ^ mice (Fig. S3C). Therefore, the lack of Tex eff-like cell differentiation may explain why Notch-deficient CD8^+^ Tex cells cannot control the infection following anti-PD-L1 treatment.

### Stromal cells expressing Delta-Like1/4 regulate the differentiation of exhausted CD8^+^ T cells

During GVHD, Tfh cell differentiation and SLEC/MPEC differentiation, SLO stromal cells expressing DLL1/4 are the major cellular source providing Notch signals to T cells^18,22,28^. Using mice where DLL1/4 genes are inactivated only in stromal cells of SLOs (CCL19-Cre+ DLL1/4^fl/fl^) or not (DLL1/4^fl/fl^), we evaluated if stromal cells were the main source of the Notch ligands to regulate CD8^+^ Tex cell differentiation. In the absence of DLL1/4 expression on stromal cells, we observed a significant increase of the TCF-1^+^ Tex-prog cell subset (Fig. 3) similar to what is observed in N1N2^Δ/Δ^ mice. The proportion of CX3CR1^+^ and CD101^+^ exhausted CD8^+^ T cells is also affected by the absence of DLL1/4 on stromal cells but to a lesser extent than what we observed in N1N2^Δ/Δ^ mice (Fig. 3). These results suggest that stromal cells of SLOs provide Notch signals to CD8^+^ T cells during chronic response, but other cell types are also likely involved.

**Figure 3.**
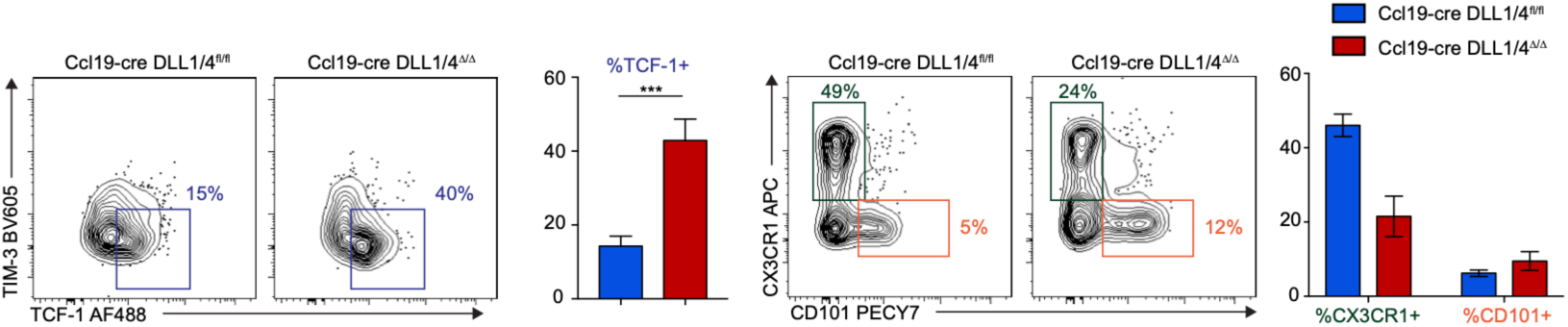
Fibroblastic reticular cells provide the Notch ligands, DLL1 and DLL4 , to prevent the severe exhaustion of CD8^+^ T cells. Control (CCL19-cre^−^ DLL1/4^fl/fl^) or mice with a specific deletion of Notch ligands DLL1 and DLL4 (CCL19-cre^+^ DLL1/4^Δ/Δ^) in stromal cells were infected with LCMV clone 13. The CD8^+^ T cell response was measured 30d later in the spleen by flow cytometry. Proportion of TCF-1^+^, CX3CR1^+^ and CD101^+^ cells was determined within gp33-specific CD8^+^ T cells in CCL19-cre^−^ DLL1/4^fl/fl^ and CCL19-cre^+^ DLL1/4^Δ/Δ^ mice is shown. Data are from two independent experiments. Unpaired Student’s *t* test, with a Welch’s correction when applied, was used for two-group comparison. \*\*\**P*<0.001.

### RNA and epigenetic profile of exhausted CD8^+^ T cells with or without Notch signaling following LCMV clone 13 infection

To investigate the role of Notch signaling in CD8^+^ T cell exhaustion, we infected Notch1/2^fl/fl^ or N1N2^Δ/Δ^ mice with LCMV clone 13 and sorted CD8⁺/CD44^hi^/PD-1⁺/Tet-GP33⁺ cells from the spleen at day 8 and 21 post-infection. Nuclei were isolated for 10x Multiome snRNA/ATAC-seq analysis(Fig. 4A).

**Figure 4.**
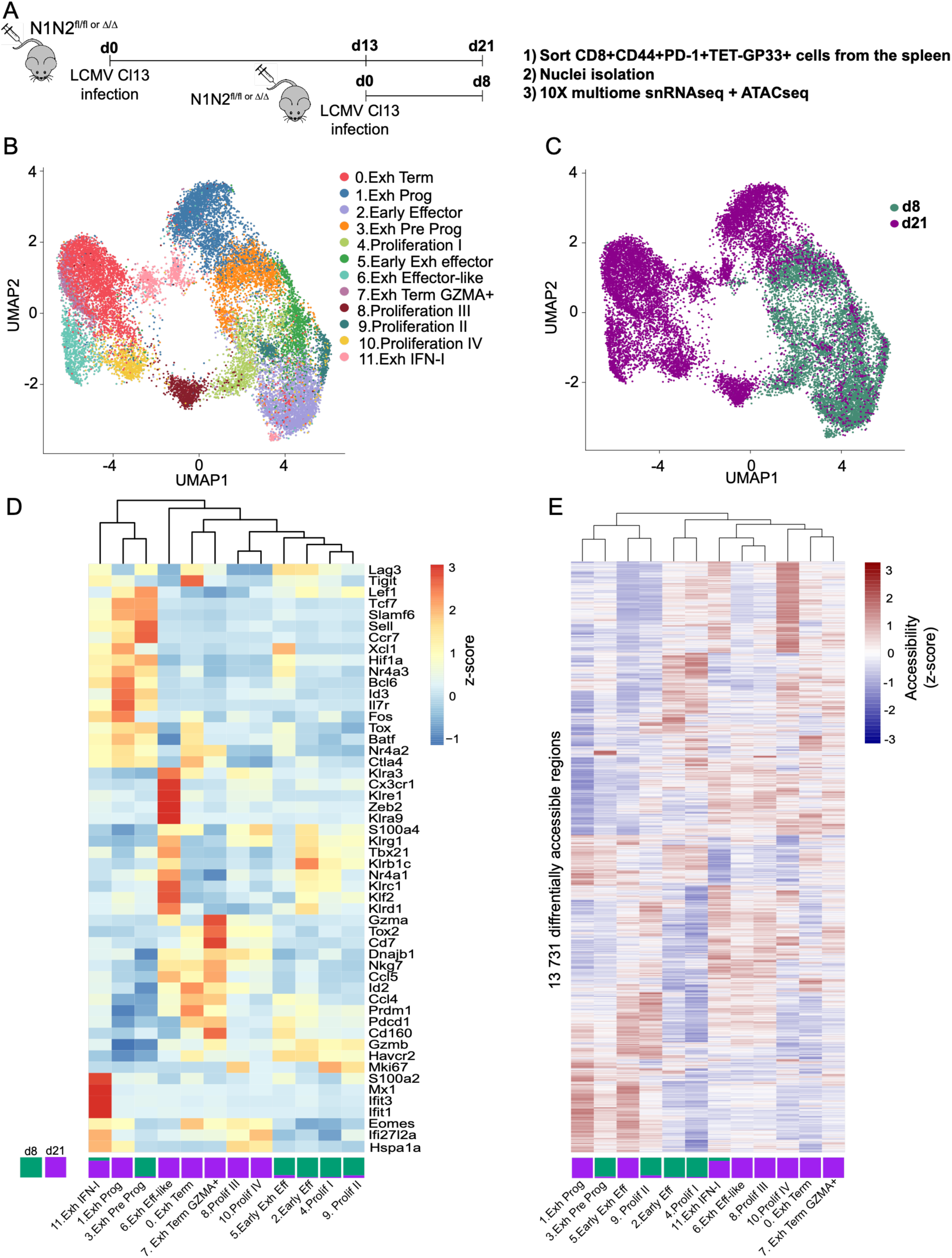
RNA and epigenetic profiles of exhausted CD8^+^ T cells with or without Notch signaling following LCMV Clone13 infection. **(A)** Experimental design. N1N2^fl/fl^ or N1N2^Δ/Δ^ mice were infected with LCMV clone 13. At days 8 and 21 post-infection, splenic CD8+CD44+PD-1+TET-GP33+ T cells were sorted, followed by nuclei isolation and Chromium 10x Genomics single-nucleus RNA and ATAC sequencing (snRNA/ATAC-seq). **(B)** UMAP visualization of integrated snRNA-seq data from day 8 and 21 post-infection, colored by transcriptionally defined CD8+ T cell subsets. **(C)** UMAP projection colored by time point day 8 versus day 21 post-infection. **(D)** Heatmap showing scaled expression (z-score) of representative marker genes across the indicated CD8+ T cell subsets, highlighting key transcriptional programs associated with each subsets. **(E)** Heatmap of differentially accessible chromatin regions (n = 13,731) across CD8+ T cell subsets, displaying normalized accessibility (z-score) and illustrating dynamic epigenetic remodeling accompanying exhaustion and differentiation states.

First, we projected all snRNA/ATAC-seq profiles into a uniform manifold approximation and projection (UMAP) space (Fig. 4B). Regardless of genotype, day-8 and day-21 cells occupied distinct areas of the UMAP, which is not surprising given the strong inflammation and abundant antigen during the acute phase compared to the chronic phase of infection (Fig. 4C). Clusters 2, 3, 4, 5, and 9 were dominant at day 8, whereas clusters 0, 1, 6, 7, 8, 10, and 11 predominated at day 21.

Next, we defined the transcriptional identity of each cluster using gene set enrichment analysis (GSEA) based on the populations described by Giles et al., *Nature Immunology* 2022 (Fig. 4D; Fig. S4A). Clusters 4, 8, 9 and 10 were enriched for proliferation-associated genes (Fig. S4B). Clusters 1 and 3 showed similar enrichment for genes associated with exhausted Progenitor cells (*Tcf7*, *Slamf6*, *Sell*, *Ccr7*). However, cluster 1 was present exclusively at day 21, whereas cluster 3 was found only at day 8. Therefore, we named cluster 1 the exhausted progenitor (Exh prog) cluster and cluster 3 the exhausted pre-progenitor (Exh Pre-prog) cluster. Clusters 0 and 7, present at day 21, exhibited similar transcriptional signatures, both corresponding to terminally exhausted T cells. A key difference between them was the expression level of *Gzma*, which was higher in cluster 7 (Fig. 4D). We therefore designated cluster 7 as Exh Term Gzma and cluster 0 as Exh Term. Cluster 2, which we refer to as early effector cells (early Eff), was found only at day 8 post-infection and was characterized by expression of *Klrg1, Tbx21,* and *Klf2* (Fig. 4D). Cluster 5 represents another effector-like population present exclusively at day 8 but distinguished by higher expression of inhibitory receptors such as *Pdcd1, Havcr2,* and *Lag3* (Fig. 4D). We therefore named cluster 5 early exhausted effector cells. Transcriptional signatures corresponding to Exh KLR population was enriched in cluster 6, which appeared only at day 21 (Fig. S4A). We designated this population as exhausted effector-like cells (Exh eff-like). This cluster was characterized by genes such as *Cx3cr1, Zeb2,* and *Klrc1* (Fig. 4D), consistent with a subset of exhausted CD8^+^ T cells retaining effector functions. Finally, cluster 11 represented a group of CD8^+^ T cells present only at day 21 whose transcriptional profile resembled type-I interferon–responsive CD8^+^ T cells (Fig. S4A). Genes such as *Mx1, Ifit1,* and *Ifit3* defined this population (Fig. 4D), which we named exhausted IFN-I–stimulated cells.

Next, we investigated the epigenetic programs established in the different Tex subsets. In total, there was 13 731 differentially accessible chromatin regions (DARs) (Fig. 4E) in which we assessed the number of DARs in each gene locus for each cluster (Fig. S5A). Using this approach, we identified global trends in DARs that were either conserved across or specific to individual Tex cell clusters. For example, Exh pre-prog and Exh prog DAR profiles were similar, including DARs at stem-associated genes, *Tcf7* and *Bach2*.

Secondly, we determined the top motif activity of transcription factors enriched in DARs that regulate transcriptional programs within each Tex subset (Fig. S5B). As shown in other studies^8,29^ , the activity of the TCF-1 and AP-1 motif was higher in Exh pre-prog and Exh prog were enriched in TCF-1 motifs. As anticipated, we also observe a higher activity of IRF motif in the exhausted IFN-I–stimulated subset. Also, Exh term Gzma had a high activity of the ETV transcription factor motifs, that was not seen in the Exh term cluster.

### Notch signaling regulates the retention and exhaustion of early Tex Prog cells

To investigate the accumulation of early Tex prog cells in the absence of Notch signaling, we first analyzed the day 8 snRNA/ATAC dataset separately. As expected, we observed a higher proportion of cells within the Exh pre-prog cluster in N1N2^Δ/Δ^ mice compared with N1N2^fl/fl^ controls (Fig. 5A). Conversely, the early effector cluster was proportionally more abundant in N1N2^fl/fl^ mice than in N1N2^Δ/Δ^ mice while the early Exh eff was more abundant in N1N2^Δ/Δ^ mice than N1N2^fl/fl^.

**Figure 5.**
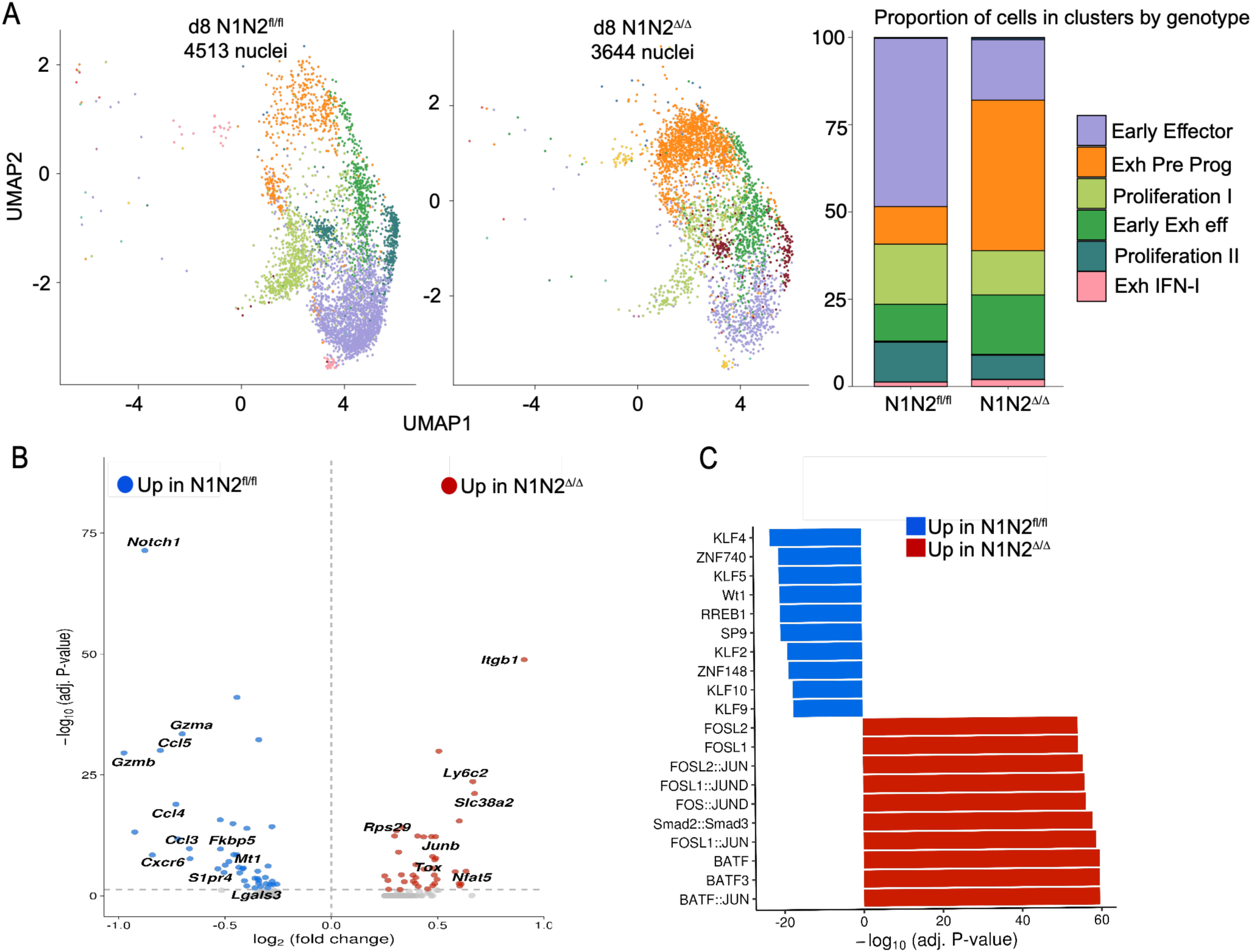
Notch signaling regulates the retention and exhaustion of early Tex prog cells. **(A)** UMAP projection of snRNA/ATAC-seq profiles from splenic CD8⁺CD44⁺PD-1⁺TET-GP33⁺ T cells isolated at day 8 post–LCMV clone 13 infection from N1N2^fl/fl^ and N1N2^Δ/Δ^ mice. The bar graph shows the proportion of cells in each cluster by genotype **(B)** Volcano plot depicting differentially expressed genes between N1N2^fl/fl^ and N1N2^Δ/Δ^ cells at day 8 post-infection in Tex pre prog (padj<0,05). Genes upregulated in N1N2^fl/fl^ cells (blue) and N1N2^Δ/Δ^ cells (red) are shown. **(C)** Transcription factor motif enrichment analysis of differentially accessible chromatin regions between N1N2^fl/fl^ and N1N2^Δ/Δ^ in Tex pre prog cells.

We focused specifically on the differentially expressed genes (DEGs) within the Tex Pre-Prog cluster in the presence or absence of Notch signaling (Fig. 5B). As expected, the transcription factor *Tox* was more highly expressed in N1N2^Δ/Δ^ cells, consistent with our flow cytometry analyses (Fig. S1C). In addition, we observed a significant upregulation of *Junb* and *Nfat5* transcripts in the absence of Notch signaling, suggesting that TCR signaling may be stronger or more sustained in N1N2^Δ/Δ^ cells compared with N1N2^fl/fl^ controls. Indeed, TCR signaling has been shown to induce *Nfat5* expression in exhausted CD8^+^ T cells during chronic LCMV clone 13 infection as well as in tumor models^30^. Moreover, *Junb* expression has been reported to be induced downstream of TCR signaling following acute infection^31^.

Recently, a TF motif enrichment analysis in DARs characterizing distinct Tex subpopulations demonstrated that BATF and AP-1 motifs are more opened in Tex pre-prog cells compared with other Tex subsets, reflecting heightened TCR signaling activity^29^. Consistent with these findings, our analyses reveal increased enrichment of BATF and AP-1 motifs in Tex pre-prog cells lacking Notch signaling compared with N1N2^fl/fl^ Tex pre-prog cells, further supporting the hypothesis that Notch-deficient exhausted CD8^+^ T cells receive stronger or more persistent TCR signals (Fig. 5C). In agreement with this, higher transcription of *Nr4a1*, *Nr4a2* and *Nr4a3*, whose expression is induced by TCR signaling^32–34^, was observed in N1N2^Δ/Δ^ Tex pre-prog cells than in their wild-type counterpart (Fig. S5C) In contrast, KLF motifs are more enriched in DARs of N1N2^fl/fl^ Tex pre-prog cells (Fig. 5C) and the KLF2 signature^35^ is lower in N1N2^Δ/Δ^ Tex pre-prog cells than in their wild-type counterpart (Fig. S6D). Notably, KLF2 has recently been shown to play a critical role in the generation of exhausted CD8^+^ T cells^36–38^. Loss of KLF2 activity induces a transcriptional signature resembling exhausted CD8^+^ T cells and impairs CD8^+^ T cell egress from SLOs, leading to their accumulation in the white pulp following LCMV clone 13 infection^39^. Furthermore, KLF2 has been shown to be necessary for the generation of Tex eff-like cells during chronic infection^37,38^. These observations further support the hypothesis that Tex pre-prog cells accumulate within SLOs and as a consequence of their inability to exit SLOs, they received persistent TCR signals in the absence of Notch signaling.

Transforming growth factor–β (TGF-β) signaling has been shown to regulate the retention of Tex prog cells within SLOs through modulation of integrin expression, notably integrins α4 (*Itga4*) and β7 (*Itgb7*)^40–42^. Expression of integrins α4 and β7 enables CD8^+^ T cells to exit SLOs and circulate, whereas TGF-β signaling suppresses α4 and β7 expression in Tex prog cells while promoting expression of integrin β1 (*Itgb1*). Among the most enriched motifs in N1N2^Δ/Δ^ Exh pre-prog cluster, we identified the Smad2::Smad3 motif, suggesting enhanced TGF-β pathway activity (Fig 5C). Binding of TGF-β to its receptors induces phosphorylation of the transcription factors SMAD2 and SMAD3, which translocate to the nucleus to activate gene transcription. Consistent with this model, *Itgb1* was the most strongly differentially expressed gene in N1N2^Δ/Δ^ Tex pre-prog cells (Fig 5B).

Collectively, our transcriptional and epigenetic analyses suggest that loss of Notch signaling promotes the retention of Tex pre-prog cells within secondary lymphoid organs, characterized by reduced chromatin accessibility at KLF2-associated loci and increased accessibility of TGF-β–associated regulatory elements. This retention likely exposes Tex pre-prog cells to heightened or prolonged TCR and TGF-β signaling, as evidenced by the significant upregulation of *Tox*, *Nfat5*, *Nr4a1*, *Nr4a2*, *Nr4a3* and *Junb* and the enrichment of DARs containing Smad motifs in the absence of Notch signaling.

### Notch signaling controls the differentiation of Tex prog into Tex eff-like cells

To understand the deficient generation of Tex eff-like cells in the absence of Notch signaling, we have deepened our analysis of the day-21 post-infection snRNA/ATAC multiome dataset (Fig. 6A). Similar to what we observed at day 8, there was a markedly higher proportion of Tex Prog cells in N1N2^Δ/Δ^ mice (34%) compared with N1N2^fl/fl^ mice (8.2%). A similar pattern was observed for the Tex Eff-like cluster, which accounted for 28% of cells in N1N2^fl/fl^ mice but only 2% in the Δ/Δ condition.

**Figure 6.**
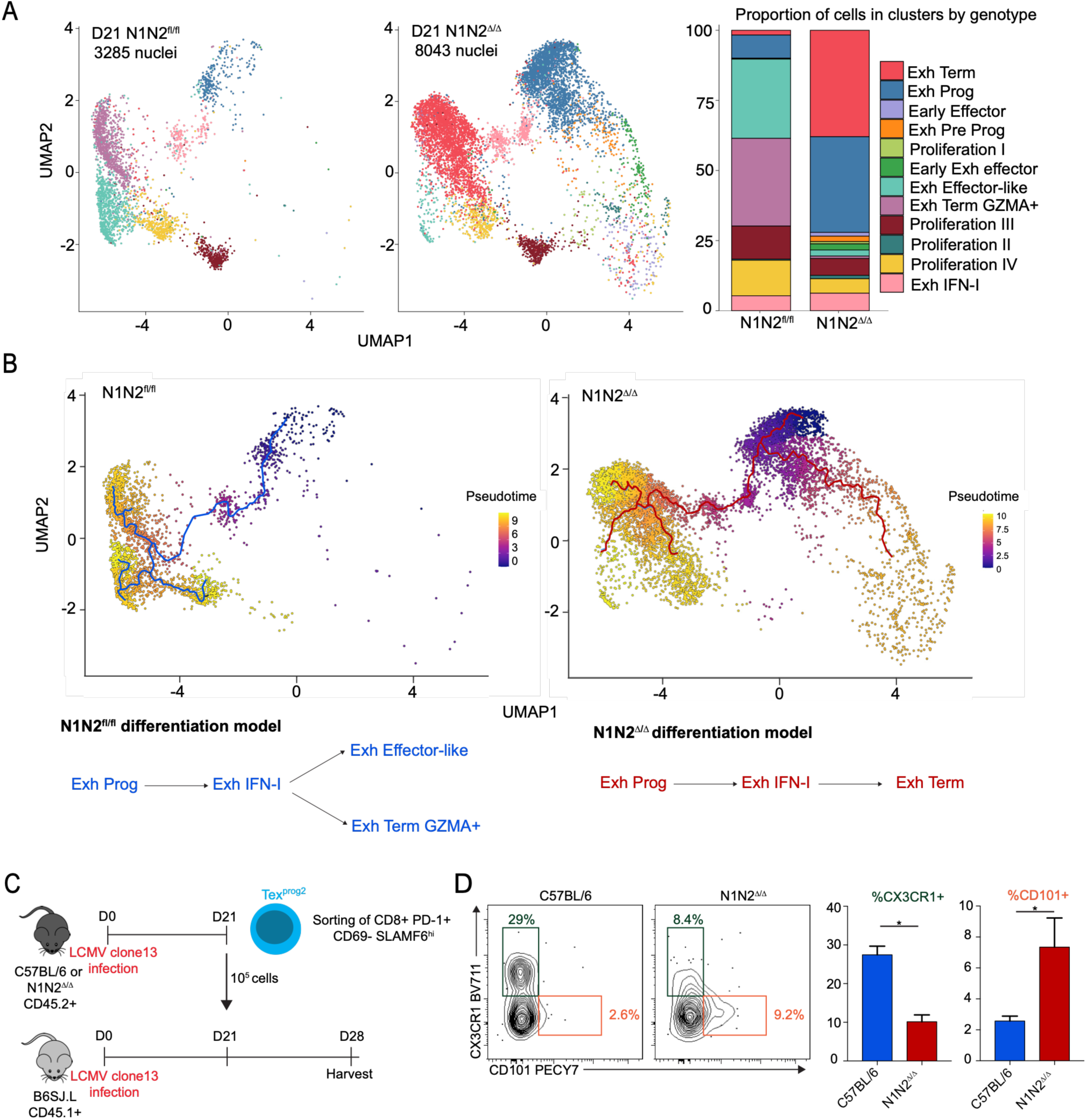
Notch signaling regulates the differentiation of CD8^+^ Tex prog cells into Tex eff-like cells. **(A)** UMAP projection of snRNA/ATAC-seq data from splenic exhausted CD8⁺ T cells isolated at day 21 post– LCMV clone 13 infection from N1N2^fl/fl^ and N1N2^Δ/Δ^ mice. Cells are colored according to transcriptionally defined clusters. The bar plot depicts the relative proportion of cells within each cluster by genotype. **(B)** Pseudotime trajectory inference overlaid onto the UMAP representation for N1N2^fl/fl^ and N1N2^Δ/Δ^ exhausted CD8⁺ T cells at day 21 post–LCMV clone 13 infection. Color gradients indicate inferred pseudotime progression, highlighting distinct differentiation trajectories in the presence or absence of Notch signaling. **(C)** Experimental design for the adoptive transfer of Tex prog2 cells (CD45.2^+^) from C57BL/6 or N1N2^Δ/Δ^ mice (day 21 post-LCMV CL13 infection) into match-infected recipients (CD45.1^+^). **(D)** The proportion of CX3CR1^+^ cells and CD101+ cells after adoptive transfer was evaluated on d7 post-transfer (gated on CD8^+^CD45.2^+^ cells). Data are representative of two experiments. Unpaired Student’s *t* test, with a Welch’s correction when applied, was used for two-group comparison \**P*<0.05.

These findings are consistent with our flow cytometry analyses (Fig. 2). However, flow cytometry did not allow us to detect the heterogeneity present within the Tex Term population. Indeed, the Tex Term cluster represented more than 30% of cells in N1N2^Δ/Δ^ mice, whereas it accounted for only 1% in N1N2^fl/fl^ mice. Conversely, the Exh Term Gzma cluster represented more than 30% of cells in N1N2^fl/fl^ mice but less than 1% in N1N2^Δ/Δ^ mice (Fig. 6A).

Our results suggest that Notch signals are important for the generation of Tex eff-like cells during chronic infection, as the absence of N1N2 receptors causes a drastic loss of CX3CR1^+^ Tex-eff cells with a concomitant increase in Tex prog1-2 cells and Tex Term cells which is also observed in the multiome analysis. Therefore, it is possible that Notch signal is necessary for the differentiation of Tex prog cells into Tex eff-like cells and that this differentiation block creates an accumulation of Tex prog cells. As we also observed an increase in Tex term cells, it is possible that Notch-deficient Tex term cells differentiate directly from Tex prog cells without going through the Tex eff-like cell state. This model is further supported by the pseudotime analysis of our mulitome (Fig. 6B). In the presence of Notch signaling, Exh prog cluster transition through the Exh IFN-I–stimulated state and then follow a bidirectional differentiation pathway, giving rise to either Exh Term Gzma or Exh Eff-like cluster. In contrast, in the absence of Notch signaling, Exh prog cluster also pass through the Exh IFN-I–stimulated state but follow a linear trajectory that leads exclusively to the Exh term population. Another interpretation could be that the Notch signal is important for maintaining the Tex eff-like cell subset following their differentiation from Tex prog cells.

To delineate how Notch signaling affects Tex cell differentiation, we sorted C57BL/6 and N1N2^Δ/Δ^ Tex prog2 cells at day 21 post-infection. These cells were adoptively transferred into LCMV clone 13 match-infected B6.SJL recipient mice (CD45.1) (Fig. 6C). At day 7 post-transfer, more C57BL/6 than N1N2^Δ/Δ^ cells have differentiated into Tex eff-like cells as indicated by the reduced expression of SLAMF6 (Fig. S6A) and the increase expression of CX3CR1 (Fig. 6D). Moreover, N1N2^Δ/Δ^ Tex prog2 cells generate significantly more Tex term cells CD101^+^ compared to their C57BL/6 counterparts (Fig. S7A and Fig 6D). Similar results were obtained after adoptive transfer of Tex prog2 into naïve B6.SJL recipients that were then infected with LCMV clone 13 (Fig. S6B-D). Thus, these results suggest that Notch is important for the differentiation of Tex prog2 cells into Tex eff-like cells and that the absence of Notch signal will promote the differentiation of Tex prog directly into Tex term cells.

These results reveal that the Notch signaling pathway is an important regulator of the differentiation of Tex prog cells into Tex eff-like cells expressing CX3CR1 and that without Notch signaling Tex prog cells differentiate into Tex term cells.

### Notch signaling regulates Tex effector differentiation by modulating the accessinility to chromatin for KLF2 transcription factors and responsiveness of CD8^+^ Tex cells to CD4 help

CD4+ T cells were shown to be necessary for the generation of Tex eff-like from Tex prog^7,13^, we performed RNAseq analysis, which provides deeper sequencing depth, on N1N2^fl/fl^ and N1N2^Δ/Δ^ gp33-specific CD8^+^ T cells at 30 post-infection with LCMV clone 13. Ingenuity pathway analysis (QIAGEN) was used to predict upstream regulators of the DEGs. Among the top 25 upstream regulators, a large proportion are cytokines and STATs, including cytokine signals involved in CD4 help to CD8^+^ T cells (Fig. 7A)^43^. Of note IL-21, the key cytokine produced by CD4^+^ T cells to promote Tex eff-like differentiation ^7^, was among the top upstream regulators (Fig. 7A). We therefore compared our transcriptomic signature with the CD4^+^ T cell help transcriptional signature identified in the context of acute infection with LCMV Armstrong^44^. ROAST analysis demonstrates that genes that are downregulated in the absence of CD4^+^ T cell help are also downregulated in the absence of Notch signaling while genes that are upregulated in the absence of CD4 help are upregulated in the absence of Notch (Fig. 7B). However, no change in IL-21 receptor transcription was observed (Fig. S7A). These results suggest that the Notch signaling pathway plays an important role to sensitize CD8^+^ T cells to cytokines and especially those involved in CD4^+^ T cell help. Thus, these results may explain the defective generation of Tex eff-like cells from Tex prog cells in absence of Notch signaling.

**Figure 7.**
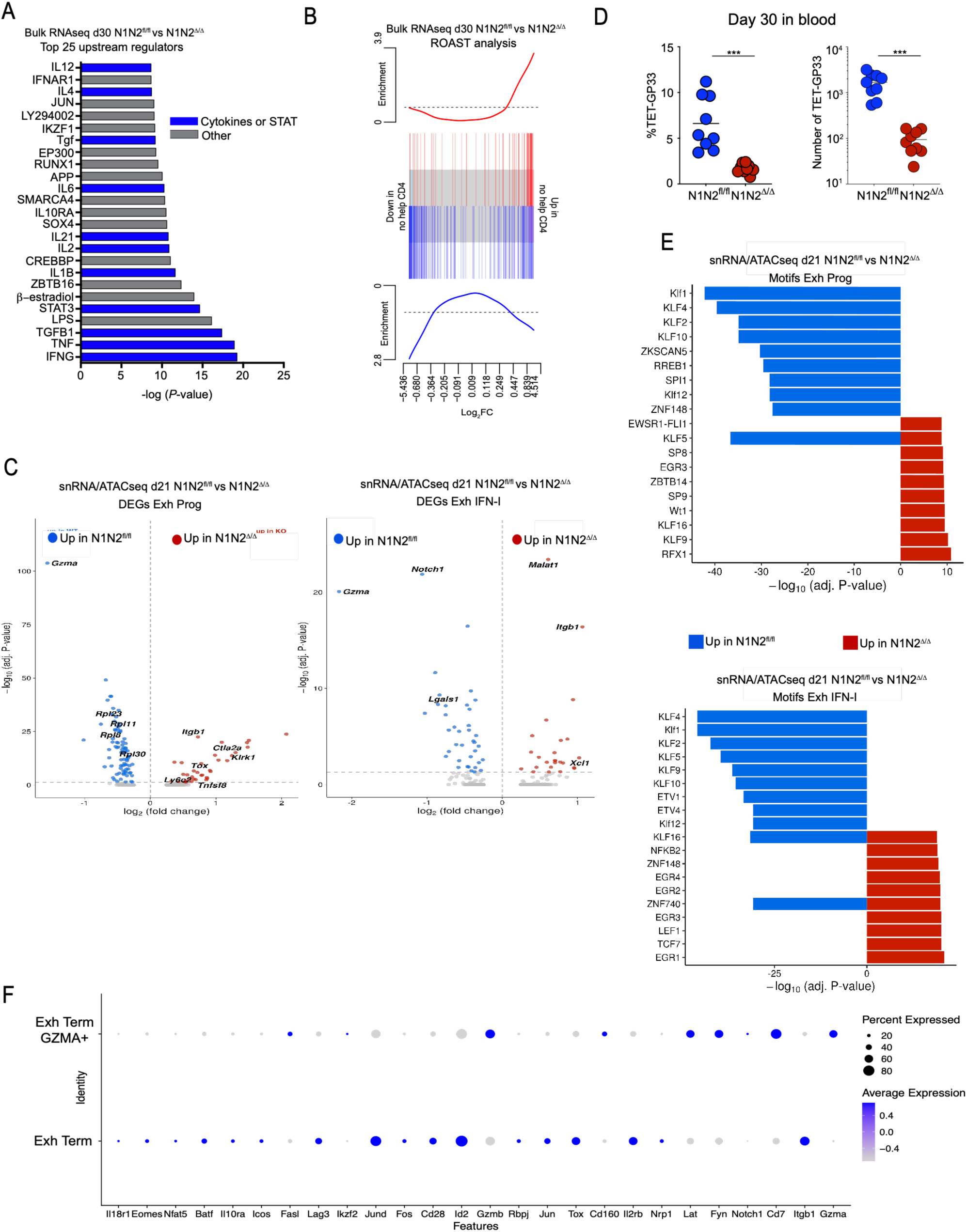
Notch signaling modulates the chromatin state and regulates de responsviness of CD8^+^ Tex cells to CD4 help. (**A)** RNAseq was performed at day 30 post infection on N1N2^fl/fl^ and N1N2^Δ/Δ^ CD8^+^CD44^+^PD-1^+^TET-gp33^+^ cells. Top 25 significant upstream regulators as determined using IPA. **(B)** ROAST enrichment analysis from gene sets from CD4-unhelped CD8^+^ T cells (Ahrends *et. al*, 2017) in the ranked lists of genes from the RNA-seq at d30. Red:genes ‘up’ in no help, blue: genes ‘down’ in no help. **(C)** Volcano plots depicting differentially expressed genes in Tex Prog (top) and Tex IFN-I (bottom) clusters from snRNA/ATAC-seq data at day 21 post-infection comparing N1N2^fl/fl^ and N1N2^Δ/Δ^ gp33-specific CD8+ T cells. **(D)** Frequency and number of gp33-specific CD8+ T cells in the blood at day 30 post-infection. **(E)** Motif enrichment analysis of differentially accessible chromatin regions in Tex Prog (top) and Tex IFN-I (bottom) clusters at day 21 post-infection. **(F)** Dot plot showing expression of selected exhaustion- and differentiation-associated genes in Tex Term and Tex Term GZMA⁺ clusters, with dot size indicating the percentage of expressing cells and color intensity representing average expression. (F) Data are representative of one experiment. Unpaired Student’s *t* test, with a Welch’s correction when applied, was used for two-group comparison \*\**P*<0.01.

Next, we compared the DEGs in the Exh prog and Exh IFN-I clusters using our snRNA/ATACseq dataset at day 21 post–LCMV clone 13 infection to identify Notch-regulated genes immediately before their bidirectional differentiation into Tex eff-like or Tex term cells (Fig. 7C). Analysis of DEGs in both Exh prog and Exh IFN-I clusters revealed increased expression of *Itgb1* and *Tox* in the absence of Notch signaling, mirroring the transcriptional profile observed in the Exh pre-prog cluster. These findings suggest that Tex prog cells lacking Notch signaling retain a more pronounced exhausted state and may reside longer within SLOs. In support of enhanced residency within SLOs and defective exit of Tex effectors, we observed a drastic reduction of LCMV-specific CD8^+^ T cells in the blood at day 30 post-infection with LCMV clone 13 (Fig. 7D). In agreement with increased residency within SLOs, within the Exh prog cluster, KLF motifs were significantly enriched in N1N2^fl/fl^, closely recapitulating the pattern observed at day 8 in Exh pre-prog cluster (Fig. 7E). Notably, these KLF motifs remained enriched during differentiation into the Exh IFN-I state (Fig. 7E). Moreover, *Klf2* transcription and chromatin accessibility at the *Klf2* locus is also decreased in absence of Notch signaling (Fig. S8B). In contrast, in the absence of Notch signaling, TCF7 motifs were enriched in the Exh IFN-I cluster, suggesting that epigenetic remodeling is delayed or impaired, as TCF7 motifs are characteristic of Tex prog cells^29^. Collectively, these results indicate that Notch signaling promotes KLF2 expression and chromatin accessibility for KLF transcription factor binding sites, thereby driving the transcriptional program required for Tex prog differentiation into Tex eff-like cells and their migration out of SLOs. Consistent with this model, enrichment of KLF motifs has been shown to be a defining feature of Tex eff-like cells^29^, and KLF2 has recently been identified as a key regulator of Tex prog to Tex eff-like differentiation^39,37,38^

Our results also indicate that Tex Prog not receiving Notch signaling will directly differentiate into Tex Term cells without going through an effector state. In absence of Notch signaling, EN1N2^Δ/Δ^ Tex term cells are transcriptionally and epigenetically distinct from their EN1N2^fl/fl^ counterpart. When comparing the expression of genes from the Exh term cluster (C0), predominantly present in N1N2^Δ/Δ^ cells, with the Exh term Gzma cluster (C7), which is mainly found in N1N2^fl/fl^ cells, a higher expression of exhaustion-associated genes can be observed in the N1N2^Δ/Δ^ Tex term (Fig 7F). This demonstrates that Notch signaling prevents CD8^+^ T cells from differentiating into severely exhausted Tex term cells. This may be the consequence of increase residency of Tex Prog in SLOs, which will induce persistent TCR and TGF-β signaling. In agreement with this, the transcription of the *Nr4a* family genes, which are induced by TCR signaling, is increased (Fig. S7C) and Smad2/3 motifs are enriched with DARs of Notch-deficient Tex term cells (Fig. S7D)

## DISCUSSION

In this study, we defined the role of the Notch signaling pathway in CD8^+^ T cells during chronic infection. We show that the Notch signaling pathway is important to prevent severe exhaustion of CD8^+^ T cell following chronic infection and acts as a master regulator of the differentiation CD8^+^ Tex subsets. We have also identified that DLL1 and DLL4 are the source of Notch ligands and demonstrated that Notch signaling is important to regulate the differentiation of CD8^+^ Tex cell subsets at different timepoint following LCMV clone 13 infection. Transcriptomic and epigenetic analyses revealed that Notch-regulated genes are linked to trafficking and cytokine response to prevent severe exhaustion and induce the differentiation of Tex prog into Tex eff-like cells.

Recent studies have described the heterogeneity of Tex cells, their different roles and sensitivity to immune checkpoint blockade. In absence of Notch signaling, we observed a significant accumulation of Tex prog and Tex term cells, but a decrease in Tex eff-like cells. Interestingly, blocking experiments with anti-DLL1/4 demonstrate a time-sensitive role for the Notch pathway on the biology of Tex subsets. Indeed, early blocking had a significant impact on the accumulation of Tex prog cells but also resulted in less Tex eff-like cells at day 30 post-infection. These results suggest that Notch signal plays an important role early in chronic infection, where permanent epigenetic modification is established^29^ and cannot be completely reversed even if the Notch signal is present at later time points.

Previous studies have demonstrated that the generation of Tex prog occurs in the early days of the response and depends on transcriptions factors TOX and TCF-1^23–26,45–48^ both of which are more expressed in absence of Notch1/2 early in the infection. TOX and TCF-1 could favor Tex Prog generation, impose an irreversible chromatin state, and inhibit other cell fates. Alternatively, it was also shown that PD-1 is an important factor to maintain TCF-1 expression ^45^ in Tex prog cells. Our results demonstrate significant increased PD-1 expression on exhausted CD8 T cells in the absence of Notch signaling, which may account for the pronounced accumulation of Tex prog via TCF-1 expression. Notch signaling could regulate negatively both transcriptions factors directly or could target TCF-1 expression by regulating PD-1 levels. Our transcriptional and epigenetic analyses suggest that Exh pre-prog cells receive enhanced TCR signaling in the absence of Notch, as evidenced by increased expression of *Nfat5*, *Tox*, *Junb* and *Nr4a* family members. These findings may partially account for the elevated PD-1 expression observed on N1N2^Δ/Δ^ Tex prog cells at day 8 post-infection, thereby contributing to their accumulation. Consistent with this model, AP-1 motifs exhibit increased chromatin accessibility in Exh pre-prog cells lacking Notch signaling, supporting the notion that exhausted CD8^+^ T cells experience stronger or more sustained TCR signaling early during the response in the absence of Notch.

Notably, our data argue that Notch signaling primarily regulates the migratory behavior of exhausted CD8 T cells rather than directly modulating TCR signal strength. Indeed, epigenetic profiling revealed decreased enrichment of KLF motifs in Exh pre-prog and Exh prog cells in the absence of Notch signaling. Recent studies have demonstrated that KLF2 promotes effector CD8 T cell differentiation by regulating cellular localisation and migration during both acute and chronic responses^36–39^.

SMAD2::SMAD3 motifs were more accessible in Exh pre-prog cells lacking Notch signaling, suggesting enhanced TGF-β pathway activity. Interestingly, TGF-β signaling has been show to control the retention of Tex prog within SLOs, via the regulation of integrin expression, and to promote their maintenance^41,42^. N1N2^Δ/Δ^ Tex prog cells exhibit increased expression of *Itgb1*, a gene regulated by TGF- β signaling and an integrin that facilitates the retention of Tex prog cells within SLOs^41,42^. This enhancement of TGF-β signaling may explain the increase in resident Tex prog that occurs without Notch signaling. Whether Notch-deficient Tex cells are per se more sensitive to TGF-β signaling or that they perceive more TGF-β within SLOs needs to be addressed. However, we believe that the failure to induce the KLF2-dependent migratory transcriptional program and the increase expression of Itgb1, which will promote SLO residency, may be responsible for the heightened sensing of TGF-β by Tex prog in absence of Notch signaling. Furthermore, the sustained TGF-β signaling in Tex prog cells may be responsible for their enhanced differentiation into Tex Term as it was shown by others that TGF-β signaling inhibits the differentiation of Tex eff while promoting the generation of Tex term cells^41,42^.

Even though there were more Tex prog cells in absence of Notch signaling, their response to anti-PD-L1 treatment was poor. This was due to the fact that they failed to differentiate into Tex eff-like cells and instead produced Tex term cells. The transition to a terminal state is also dictated by important changes in chromatin accessibility, which could be regulated by Notch^8,49^. These results suggest that Notch plays important roles at different time points of the chronic infection.

Two studies suggested that the differentiation model of Tex subsets is linear, where Tex prog cells sequentially give rise to Tex eff-like cells and Tex term cells^8,9^, but another study proposed that Tex prog can give rise to both Tex eff-like or Tex term^7^. Our results are in line with the latter model, where the absence of Notch signals will cause the differentiation of Tex prog into Tex term without transiting through a Tex eff-like state. Indeed, adoptive transfer of Tex prog lacking Notch1/2 expression showed that these preferentially differentiate into Tex-term cells. However, it is also possible that, in absence of Notch signaling, Tex eff-like rapidly differentiate into Tex term, although this is not supported by experiments where we adoptively transferred WT Tex eff-like into mice treated with anti-DLL1/4.

Pseudotime analysis of our snRNA/ATACseq dataset at day 21 post–LCMV clone 13 infection revealed that exhausted progenitor (Exh prog) CD8^+^ T cells initially differentiate into an interferon-responsive exhausted population (Exh IFN-I), which subsequently bifurcates into either Tex eff-like or Tex term states supporting the results obtained with the adoptive transfer of the Tex prog. In contrast, loss of Notch signaling enforces a linear differentiation trajectory in which Exh IFN-I cells exclusively progress toward a terminally exhausted fate. Notably, Tex term cells generated in the absence of Notch signaling exhibit a more severe exhaustion phenotype, characterized by reduced *Gzma* expression compared with Tex term cells from N1N2^fl/fl^ controls. Consistent with a previous report identifying an IFN-I–sensitive intermediate population driving Tex term differentiation^14^, transcriptional profiling of bulk RNAseq data at day 30 post-infection demonstrated enrichment of an IFN-I response signature in Notch-deficient CD8^+^ T cells suggesting that heightened IFN-I signaling might be driving the differentiation of Tex prog into Tex term in the absence of Notch signaling.

Epigenetic analyses further revealed increased accessibility of KLF motifs in Exh prog cells in the presence of Notch at day 21 post-infection, mirroring observations at day 8 and suggesting sustained KLF transcription factors activity in N1N2^fl/fl^ Tex prog cells. Consistent with this, KLF2 has been shown to restrain Tex Prog–to–Tex Term differentiation in both tumor and chronic LCMV infection models, while loss of KLF2 activity is associated with enhanced IFN-I–driven transcriptional program and reduces generation of Tex eff-like cells^36–39^. Together, these findings suggest a model in which Notch signaling limits IFN-I–mediated terminal exhaustion by maintaining KLF2-dependent chromatin accessibility and migratory programs in Tex prog cells. Whether Notch primarily regulates intrinsic sensitivity to IFN-I or instead restricts IFN-I exposure by promoting KLF2-dependent egress from secondary lymphoid organs remains an important question for future investigation. Alternatively, Notch signaling itself might be required for the proper transcriptional induction of *Klf2* within Tex cells.

The transcriptomic analyses of our bulk RNAseq at day 30 post-LCMV clone 13 infection, suggested that the Notch signaling pathway plays an important role in sensitizing CD8^+^ T cells to cytokine signals. Indeed, the top 25 of the upstream regulators in absence of Notch are mainly cytokines and STATs, including IL-21 and IL-2. Moreover, most of these cytokines are provided in the context of CD4^+^ T cell help^43^. CD4^+^ T cell help was demonstrated to be important for Tex cell differentiation. Indeed, recent studies showed that the differentiation of Tex prog into Tex eff-like cells is dependent on CD4^+^ T cells help via IL-21^7,13,50^. Notch signals could be important to allow CD8^+^ T cells to respond properly to the cytokines generated by CD4^+^ T cells that will then promote Tex prog differentiation into Tex eff-like. In agreement with this possibility, we detected an enrichment of the unhelped signature in absence of Notch signaling in our RNAseq data. The effect of Notch signaling on the transcriptional response to cytokine stimulation is reminiscent of what we have observed during acute infection^18^.

Finaly, while we demonstrated that stromal cells expressing Notch ligands DLL1/4 are an important source of the Notch signal during chronic infection, other cell types, such as antigen presenting cells and endothelial cells, seem to provide Notch signal to modulate CD8^+^ T cell exhaustion. Indeed, the results showed that the absence of ligands DLL1/4 on stromal cells only partially recapitulate phenotypes observed in CD8^+^ T cells deficient for Notch1/2. In acute infection and GVHD, stromal cells expressing DLL1/4 are the almost exclusive source of Notch signal that modulate SLEC differentiation and T cell function, respectively^18,51^. It was shown that the stromal cells are an important target of LCMV clone 13 and their functions are altered^52^, but the impact of infection on their ability to provide Notch signal has not been evaluated. Perhaps destruction of stromal architecture that occurs in chronic infection results in APCs becoming more important in supplying different signals, including Notch ligands. Alternatively, persistent inflammation may drive APCs expression of DLL1/4.

Together, our results suggest that the Notch signaling pathway could be modulated to tune the populations in chronic settings. When activated, it could offer the possibility to temporarily increase the number of effectors, when needed. Alternatively, Notch-inhibition could be used to favor an accumulation of progenitors, during adoptive cell therapy, for example. For these reasons, targeting Notch is a promising therapeutic strategy in chronic infection or cancer.

## MATERIALS AND METHODS

### Mice

B6.SJL, C57BL/6 and mice were bred at the Maisonneuve-Rosemont Hospital Research Center facility. Notch1^fl/fl^/Notch2^fl/fl^ (N1N2^fl/fl^) and E8I-cre^+/-^ Notch1^fl/fl^Notch2^fl/fl^ (N1N2^Δ/Δ^)^16^, Notch1^fl/fl^/Notch2^fl/fl^ P14 mice and E8I-cre^+/-^ Notch1^fl/fl^Notch2^fl/fl^ P14 mice were generated by crossing a P14 transgenic mice with Notch1^fl/fl^/Notch2^fl/fl^ (P14N1N2^fl/fl^) or E8I-cre^+/-^ Notch1^fl/fl^Notch2^fl/fl^ (P14N1N2^Δ/Δ^). Mice were bred at the Maisonneuve-Rosemont Hospital Research Center facility and housed in a pathogen-free environment and treated in accordance to the Canadian Council on Animal Care guidelines. Following infection, mice were monitored daily for weight loss, dehydration and lethargy. Our animal protocol was approved by the Hospital Maisonneuve-Rosemont Council on Animal Care. Delta-like1^fl/fl^Delta-like4^fl/fl^, Ccl19-cre^+/-^ Delta-like1^fl/fl^Delta-like4^fl/fl^ mice were previously described in^22,53^ and were bred at the University of Pennsylvania. All the animal protocol for these mice was approved by the University of Pennsylvania’s Office of Regulatory Affairs and Care of Animals.

### Adoptive cell transfer of P14N1N2 fl/fl or Δ/Δ CD8^+^ T cells

Naive P14N1N2^fl/fl^ or P14N1N2^Δ/Δ^ CD8+ T cells were isolated from lymph nodes using the Stemcell naive CD8+ T cells isolation kit and 2 × 10^3^ cells were adoptively transfer by intravenous (i.v.) injection in recipient mice. 18-24 hours later, mice were infected with LCMV clone 13 as described below.

### LCMV clone 13 infection

The virus LCMV clone 13 was produced by infection of the L929 fibroblast cell line, cultured in Minimum Essential Medium (MEM) containing 5% heat inactivated Nu serum, followed by harvesting of the produced virus in the supernatant. The virus titer was determined using MC57G fibroblasts as previously described^54^. Mice were infected with 2 × 10^6^ PFUs of LCMV clone 13 by i.v. injection.

### Anti-PD-L1 blocking antibodies treatment

Following LCMV clone 13 infection, N1N2^fl/fl^ or N1N2^Δ/Δ^ mice were injected i.p. with 0.2 mg of anti-mouse PD-L1 (clone: 10F.9G2) or the isotype control from BioXcell every 3 days between day 23 and day 35 post-infection. Mice were sacrifice 37 days post-infection.

### Anti-IFNAR blocking antibodies treatment

C57Bl/6 mice were treated i.p. with IFNR1 blocking antibody (clone MAR1-5A3; Leinco Technologies, St. Louis MO) or a mouse IgG1 isotype control (clone MOPC21) either starting prior to infection: day -1 (500µg), day 0 (500µg), day 3 (250µg) and day 6 (250µg).

### Adoptive transfer of Tex cell subsets

First, donors (CD45.2) C57BL/6 or N1N2^Δ/Δ^ mice were infected with LCMV clone 13. At the same time recipient (CD45.1) B6.SJL were infected with LCMV clone 13. At day 21 post-infection, donor mice were sacrificed and the spleens were collected. Before cell sorting, CD4 T cells and CD19+ B cells were removed from the cell suspension using the StemCell rapidsphere strepatividin kit. After, the cell suspension was stained with a Zombie dye and the following markers: anti-CD8, anti-CD44, anti-PD-1, anti-CD69 and anti-SLAMF6. Tex^prog2^ cells (Zombie^neg^, CD8+, CD44+, PD-1+, SLAMF6^hi^, CD69-) or Tex^eff^ (Zombie^neg^, CD8+, CD44+, PD-1+, SLAMF6^lo^, CD69-) were sorted with a BD FACSARIA III and adoptively transfered (10^5^ Tex^prog2^ cells or 7 × 10^4^ Tex^eff^ cells) in match-infected mice or naïve mice, depending of the experiment. Recipient mice were sacrificed 7 or 8 days later.

### Cell preparation, antibodies, flow cytometry and cell sorting

Spleens were dissociated using frosted glass slides. Red blood cell lysis was performed on spleens using 0.83% NH_4_Cl for 5 min at room temperature (RT). Tetramer H2D^b^-gp33_33-41_ (NIH) staining was performed at 37°C for 15 min. Extracellular staining was performed for 20 min at 4°C as previously described^55^. Intranuclear staining for transcription factors was performed using the Foxp3/Transcription Factor Staining Buffer Set according to manufacturer (Invitrogen) instructions. A complete list of antibodies used in this article is available in Table S3. Flow cytometry analysis was performed on a BD LSRFortessa X-20 from BD Biosciences and data were analyzed using FlowJo software (BD).

Cell sorting for RNAseq of N1N2^fl/fl^ or ^Δ/Δ^ cells was performed on ex vivo splenocytes. Briefly, cell suspensions were incubated with viability dye and Fc-block antibodies for 10 min at RT. Cell suspensions were then stained with extracellular antibodies 20 min at 4 °C in sorting buffer (PBS, 1% Nu serum, 1 mM EDTA, 25 mM Hepes). Viable cells (Zombiedye^−^CD8^+^ CD45.2^+^) were then sorted with a BD FACSAria III.

### Ex vivo restimulation of CD8^+^ T cells with peptide gp33 for analysis of cytokine production

Ex vivo splenocytes were collected and stimulated with the gp33 peptide (0.1 μg/mL) in the presence of brefeldin A (BFA, 10 μg/mL) for 5 h at 37 °C. Following stimulation, cells were fixed, permeabilized, and stained as previously described ^56^. Briefly, restimulated cells were fixed 20 min with 1% paraformaldehyde at RT. Fixed cells were permeabilized with saponin (0.5%) for 10 min at RT. Permeabilized cells were stained with anti-cytokine antibodies followed by cell surface staining.

### RNAseq sample preparation

First, N1N2^fl/fl^ or N1N2^Δ/Δ^ mice were infected with LCMV clone 13. At day 30 postinfection, exhausted (CD8^+^CD44^+^PD-1^+^TET-GP33^+^) N1N2^fl/fl^ or N1N2^Δ/Δ^ cells were sorted directly into TRIzol reagent. For day 8 and day 30 post-infection, a total of 6 individual biological samples (3 mice per group of two independent experiments) were collected for each genotype and time point (3 × 10^4^ to 7.5 × 10^4^ cells per sample). RNA extraction, library preparation and sequencing was done at the IRIC Genomics Platform (University of Montreal). The cDNA library sequencing was performed on Illumina Nextseq500 (75 cycles single-end reads).

### RNAseq analysis

Bioinformatical analyses were done at the IRIC Genomics Platform and at the Institut de recherches cliniques de Montréal (IRCM). Sequenced reads were trimmed for sequencing adapters and low-quality 3ʹ bases using Trimmomatic version 0.35^57^ and then aligned to the reference mouse genome version GRCm38 (or mm10 gene annotation from Gencode version M13, based on Ensembl 88) using STAR version 2.5.1b^58^. Gene expressions were obtained both as readcount directly from STAR as well as computed using RSEM^59^ to obtain gene and transcript level expression, either in transcripts per kilobase million or fragments per kilo base per million mapped reads values, for these stranded RNA libraries. DESeq2 version 1.16.1^60^ was then used to normalize gene readcounts and compute differential expression between the different experimental conditions. Determination of relevant upstream regulators was done with QIAGEN’s Ingenuity Pathway Analysis (IPA).

For ROAST enrichment analysis^61^, the differential expression analysis of the RNAseq above was performed with the edgeR^62^ and limma^63^ packages in R. To allow comparisons of different samples, TMM normalization (relative to the total number of reads per sample) and VOOM transformation were applied^64^.Only genes with log2 counts greater than 0.5 counts per million reads in at least 2 of the samples were kept for further analysis. The linear model was then fitted with the lmFit function. Differential expression was assessed using empirical Bayes moderated t-statistics with the eBayes function while controlling the false discovery rate (FDR) at 5%. The gene set used as a reference for the analysis is issued from^44^, accessible through accession number GSE89665 in the supplementary file GSE89665 COMP3 DE Table.txt.gz. Differentially expressed genes for the gene set were determined as those with a false discovery rate less than 5% and absolute logFoldChange value greater than 2. ROAST tests were performed with the roast function of the limma package, using the ‘mean’ set statistic and 10000 rotations.

Enrichment for the IFN-I response genes was tested using publicly available mouse data (GSEA MM3877). Mouse data from GSEA MM3877 were imported and analyzed using Seurat.

### Nuclei isolation for snRNA/ATACseq

Isolation of splenic CD8 T cells was executed as described previously. Cell sorting was performed as described previously, except the cells were sorted 2 times to obtain the most pur populations. Then, purified CD8+ T cells were processed immediately for nuclei isolation following the Chromium Next GEM Single Cell Multiome ATAC + Gene Expression protocol (10x Genomics). Briefly, cells were resuspended in chilled lysis buffer optimized for lymphocytes and incubated on ice to selectively lyse the plasma membrane while preserving nuclear integrity. Nuclei were washed, filtered through a 40-µm strainer, and resuspended in nuclei buffer. Nuclei concentration and integrity were assessed using trypan blue staining and light microscopy.

### Chromium Next GEM Multiome library preparation

Isolated nuclei were loaded onto the Chromium Next GEM chip targeting 5,000–10,000 nuclei per sample. Gel bead–in–emulsion (GEM) generation and barcoding were performed using the Chromium Controller. Within individual GEMs, transposition of accessible chromatin regions was carried out simultaneously with reverse transcription of nuclear mRNA, allowing the capture of chromatin accessibility and gene expression profiles from the same nucleus.

Following GEM recovery, barcoded ATAC and gene expression libraries were amplified separately according to the manufacturer’s protocol. ATAC libraries were size-selected to enrich for nucleosome-free and mono-nucleosomal fragments, whereas gene expression libraries were amplified to capture full-length cDNA.

### Sequencing

Final libraries were quantified using Qubit and fragment size distributions were assessed on Bioanalyzer. Libraries were sequenced on an Illumina NovaSeq 6000 platform. Gene expression libraries were sequenced with a minimum depth of 50,000 read pairs per nucleus, and ATAC libraries at a minimum depth of 40,000 read pairs per nucleus.

### Multiome data processing and analysis

Joint RNAseq and ATACseq single-cell multiome data were generated using the 10x Genomics Chromium platform for four samples. Raw sequencing data were processed using Cell Ranger ARC (v2.0.0, 10x Genomics) and aligned to the mouse GRCm38 reference genome to generate gene expression and chromatin accessibility count matrices.

Downstream analyses were performed in R using the Seurat (v4.1.3) and Signac (v10.0) packages. Each sample was initially processed independently. Cells were first filtered based on standard RNA-seq quality control metrics (500 < nFeature_RNA < 4000 & nCount_RNA < 18000 & percent.mt < 30), followed by filtering based on ATAC-seq quality metrics (3000 < nCount_ATAC < 80000 & nucleosome_signal < 1.2 & TSS.enrichment > 1), to remove low-quality cells.

Following quality control, filtered gene expression data were normalized using log-normalization and highly variable genes identified. Counts were scaled with the cell cycle variation regressed out and dimensionality reduction was performed using principal component analysis (PCA). The first 30 PCs were used for downstream analysis. For the filtered chromatin accessibility data, ATAC-seq counts were normalized using term frequency–inverse document frequency (TF-IDF) transformation, followed by dimensionality reduction using latent semantic indexing (LSI). All samples were integrated across batches separately for the RNA-seq and ATAC-seq assays using Seurat and Signac integration workflows. The integrated RNA and ATAC assays were subsequently combined using weighted nearest neighbor (WNN) analysis, enabling joint representation of transcriptional and chromatin accessibility information. The resulting multi-modal object was processed following recommended Signac workflows, including dimensionality reduction and graph-based clustering.

Cluster-specific marker genes were identified from the RNA-seq assay using differential expression analysis. These markers were used to manually annotate the clusters found. Differentially accessible regions (DARs) were identified for each cluster from the ATAC-seq assay. Motif enrichment analysis of DARs was performed using transcription factor motifs from the JASPAR database, and chromatin accessibility deviations were quantified using chromVAR, as implemented in Signac. Differential motif activity was assessed across clusters to identify candidate transcription factors associated with distinct cell states.

Finally, pseudotime analysis was performed independently for each sample using Monocle3 to infer potential cellular trajectories.

All visualizations and statistical analyses were performed in R.

### Statistical Analysis

Statistical analyses for differences between the fl/fl and Δ/Δ groups were done using Student’s T test. Welch’s correction was applied for unequal variances when required. ANOVA was used when comparing more than two experimental groups. Tukey’s correction was applied for unequal variances when required. Data are presented as mean +/- standard error of the mean (SEM). Only significant statistical differences are indicated on the figures.

## Data availability

Data will be deposited in a publicly available database.

## Author contributions

DMDS, EP, FD, MEL and JFD performed the experiments. DMDS, SB and LLC analyzed the data. DMDS, SB, IM and NL prepared the manuscript. AL, CS, FR and BL provided critical reagents. IM and NL supervised the work.

## Acknowledgements

We would like to acknowledge all lab members for helpful discussion. M. Dupuis for cell sorting and animal care technicians for animal husbandry. We are grateful to V. Calderon and C. Grou from IRCM for bioinformatical analysis. This work was supported by a grant from the Canadian Institutes of Health Research (PJT-152988) to N.L. and from the National Institute of Allergy and Infectious Diseases (NIAID; R01-AI091627) to I.M. EP was supported by T32-GM007863 from the National Institute for General Medical Sciences (NIGMS) and F30-AI136325 from NIAID. DMDS and LLC were supported by a studentship from the Fonds de la Recherche Québec-Santé.

## Abbreviations

Ag: antigen
APC: antigen presenting cell
BFA: Brefeldin A
Exh: Exhausted
DAR: Differentially accessible regions
DEG: differentially expressed gene
DLL: Delta-like
eff-like: effector-like
FFU: Focus forming units
gp33: glycoprotein 33
GzmB: Granzyme B
IR: inhibitory receptor
GVHD: graft-versus-host disease
IPA: Ingenuity Pathway Analysis
LCMV: Lymphocytic choriomeningitis virus
mAbs: monoclonal antibodies
MPEC: memory precursor effector cell
NICD: notch intracellular domain
N1: Notch1
N2: Notch2
prog: progenitor
RT: room temperature
SLEC: short-lived effector cell
SLO: secondary lymphoid organ
term: terminal
Tex: T exhausted
WT: wild-type

