## Supplemental Figures for "The Notch signaling pathway is a master regulator of CD8^+^ T cell exhaustion and differentiation during chronic infection"

#### SUPPLEMENTARY DATA

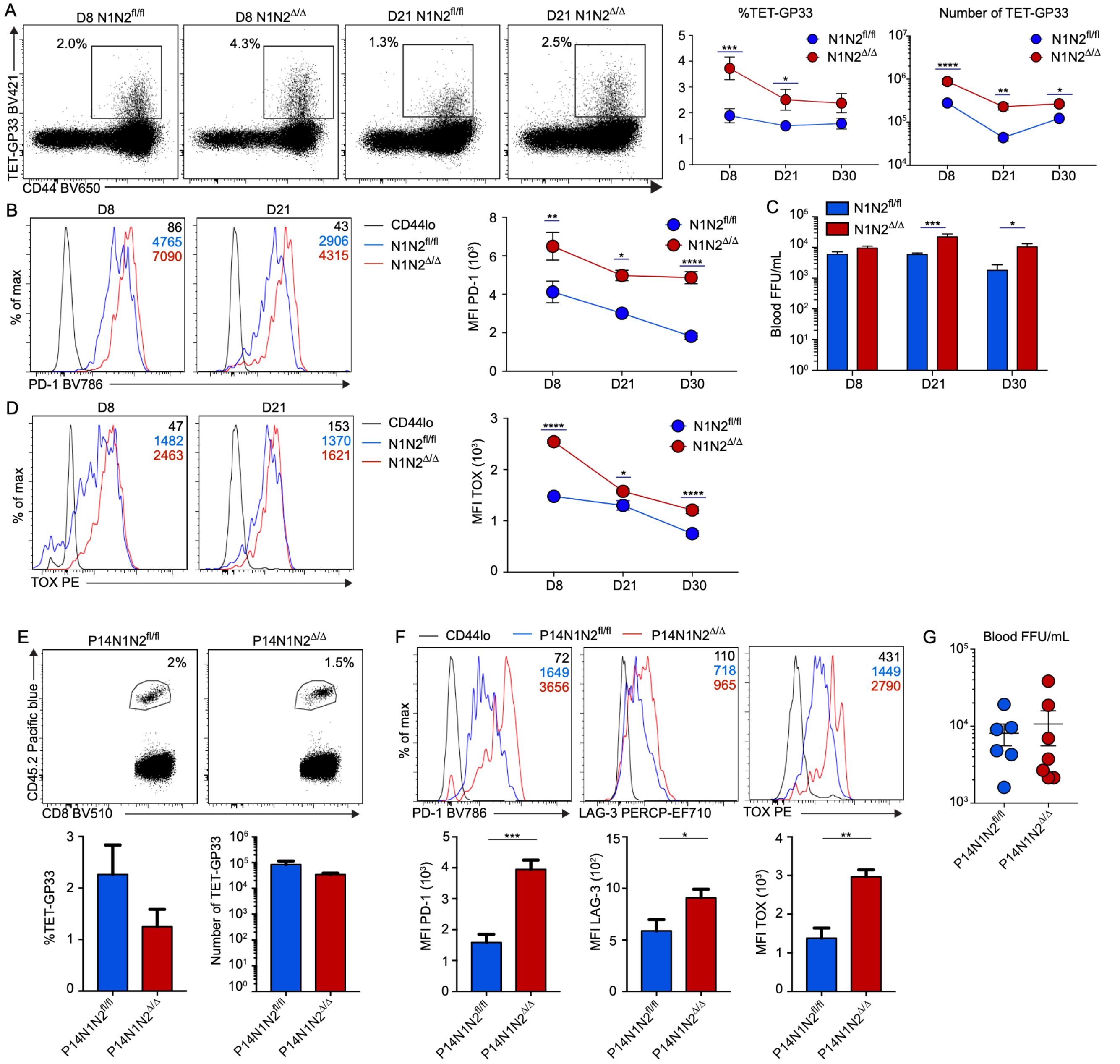


**Figure S1. The Notch signaling pathway prevents severe CD8⁺ T cell exhaustion throughout the infection independently of viral load following LCMV clone 13 infection.**

**(A-C)** Mice were infected with LCMV clone 13 and the response in the spleen was characterized at d8, d21 and d30 post-infection by flow cytometry. (**A**) Percentage and numbers of gp33-specific CD8⁺ T cells. Expression of inhibitory receptor PD-1 **(B)** and transcription factor TOX (**C**) on gp33-specific N1N2$^{fl/fl}$ mice and N1N2$^{\Delta/\Delta}$ CD8⁺ T cells over the course of the infection. (**D**) Viral load in the blood was determined using focus-forming assay on MC57G cells. **(E-G)** Adoptive transfer of P14N1N2$^{fl/fl}$ or P14N1N2$^{\Delta/\Delta}$ cells (CD45.2⁺) into B6.SJL recipients 1d before the infection with LCMV clone 13. The response in the spleen and the viral load were characterized at d30 post-infection by flow cytometry. (**E**) Percentage and numbers of CD45.2$^{+}$ CD8$^{+}$ T cells P14N1N2$^{fl/fl}$ and P14N1N2$^{\Delta/\Delta}$ cells. (**F**) Expression of inhibitory receptors (PD-1 and LAG-3) and transcription factor TOX was measured on CD8$^{+}$CD45.2$^{+}$ P14N1N2$^{fl/fl}$ and P14N1N2$^{\Delta/\Delta}$ cells. (**G**) Viral load in blood**.** Data are from 2 independent experiment (A-C) and data are representative of 1 from 2 independent experiments (D-F). Unpaired Student's *t* test, with a Welch's correction when applied, was used for two-group comparison. **P*<0.05, ***P*<0.01, ****P*<0.001, *****P*<0.0001.

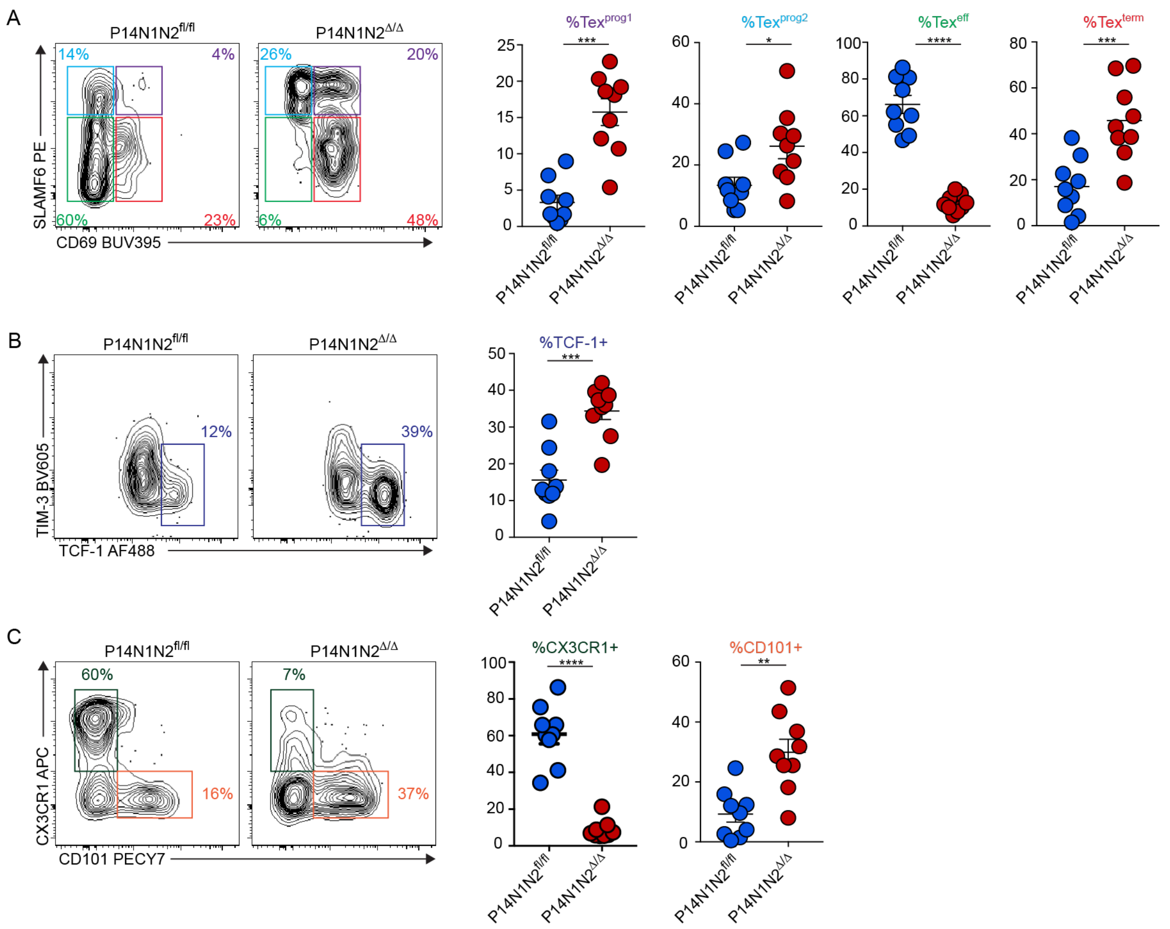


**Figure S2. Notch signaling acts as a master regulator of exhausted CD8$^+$ T cell differentiation even when the viral loads are normalized.**

Adoptive transfer of P14N1N2$^{fl/fl}$ or P14N1N2$^{\Delta/\Delta}$ cells (CD45.2$^+$) into B6.SJL mice was done 1d before LCMV clone 13 infection. The response in the spleen was characterized at d30 post-infection using flow cytometry. Proportion of Tex subsets **(A),** TCF-1$^+$ cells (**B**) and CX3CR1$^+$ or CD101$^+$ cells (**C**) amongst CD45.2$^+$ P14N1N2$^{fl/fl}$ or P14N1N2$^{\Delta/\Delta}$ cells. Data are from two independent experiments. Unpaired Student's *t* test, with a Welch's correction when applied, was used for two-group comparison. *$P$<0.05, **$P$<0.01, ***$P$<0.001, ****$P$<0.0001.

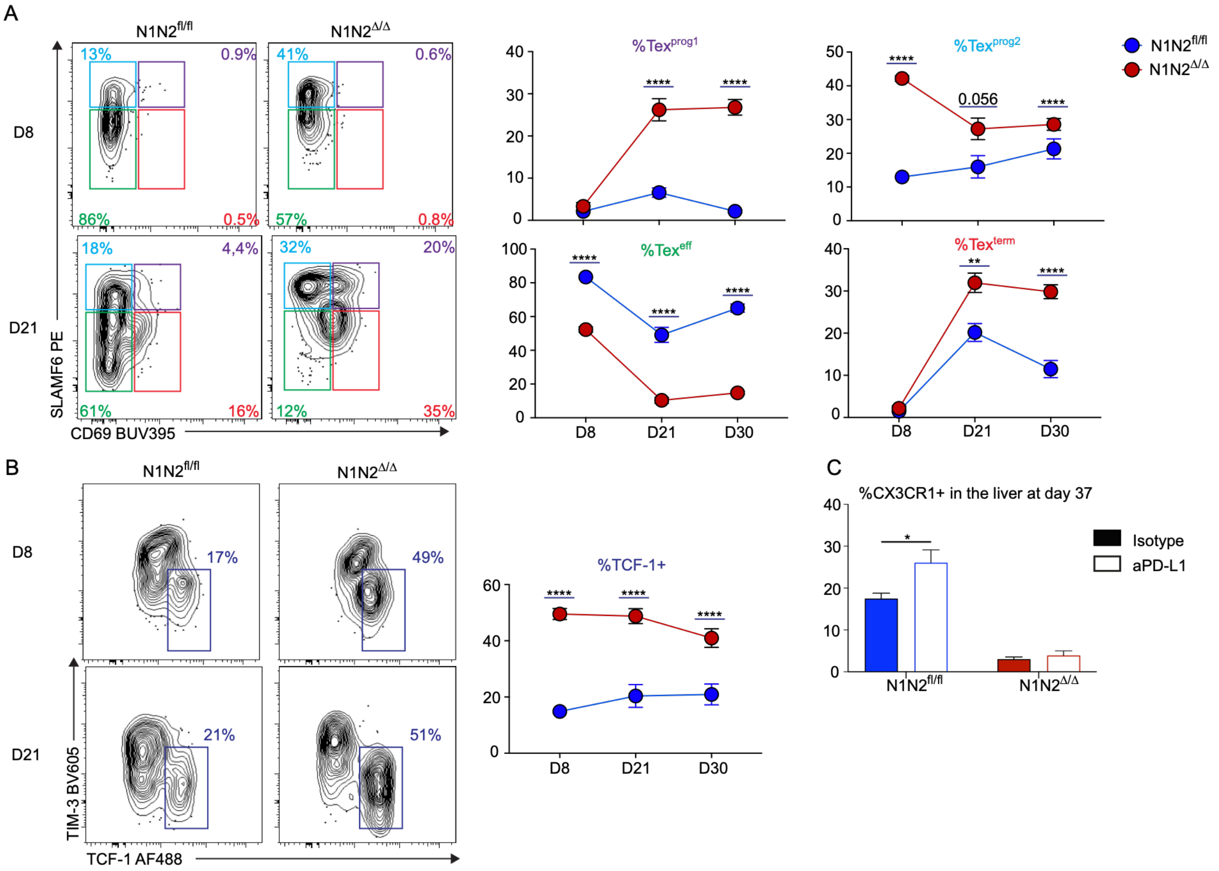


**Figure S3. Notch signaling acts as a master regulator of exhausted CD8⁺ T cell subset differentiation throughout LCMV clone 13 infection.**

Mice were infected with LCMV clone 13 and the TET-gp33⁺ response in the spleen was characterized at different time points by flow cytometry. **(A)** Proportion of Tex subsets within gp33-specific CD8⁺ T cells from N1N2$^{fl/fl}$ and N1N2$^{\Delta/\Delta}$ mice. (**B**) Proportion and numbers of TCF-1+ cells on gp33-specific CD8⁺ T cells from N1N2$^{fl/fl}$ and N1N2$^{\Delta/\Delta}$ mice. **(C)** Anti-PD-L1 treatment does not increase the differentiation into Tex eff-like cells in absence of Notch signaling. N1N2$^{fl/fl}$ and N1N2$^{\Delta/\Delta}$ mice were infected with LCMV clone 13 and treated with 0.2 mg of anti-PD-L1 or isotype control every 3 days between day 23 and day 35 post-infection. Proportion of Tex eff-like cells (CX3CR1⁺) in the liver were determined on gp33-specific CD8⁺ T cells from N1N2$^{fl/fl}$ and N1N2$^{\Delta/\Delta}$ mice at day 37 post-infection. Data are from two independent experiments. Unpaired Student's *t* test, with a Welch's correction when applied, was used for two-group comparison. $^{**}P<0.01$, $^{****}P<0.0001$.

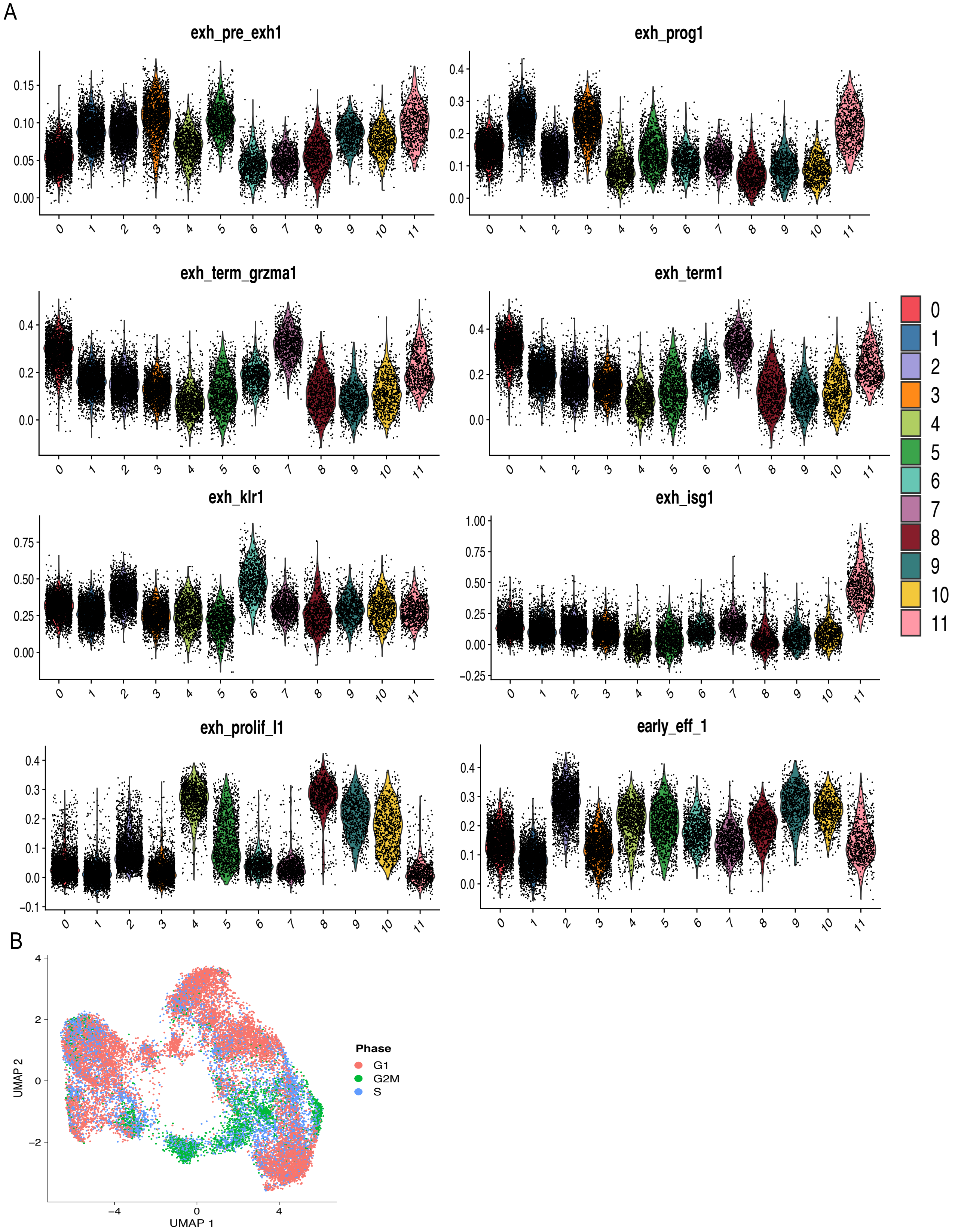


**Figure S4. Gene set enrichment analysis and cell cycle status across exhausted CD8⁺ T cell subsets at day 8 and day 21 post–LCMV clone 13 infection.**

**(A)** Violon plot showing the enrichment of gene signature defining CD8 Tex subsets (described by Giles et al., *Nature Immunology* 2022) within the different clusters of our snRNA/ATACseq. Each dot represents an individual nucleus, colored according to cluster identity.
**(B)** UMAP projection of all nuclei colored by inferred cell cycle phase (G1, S, or G2/M) using mRNA data.

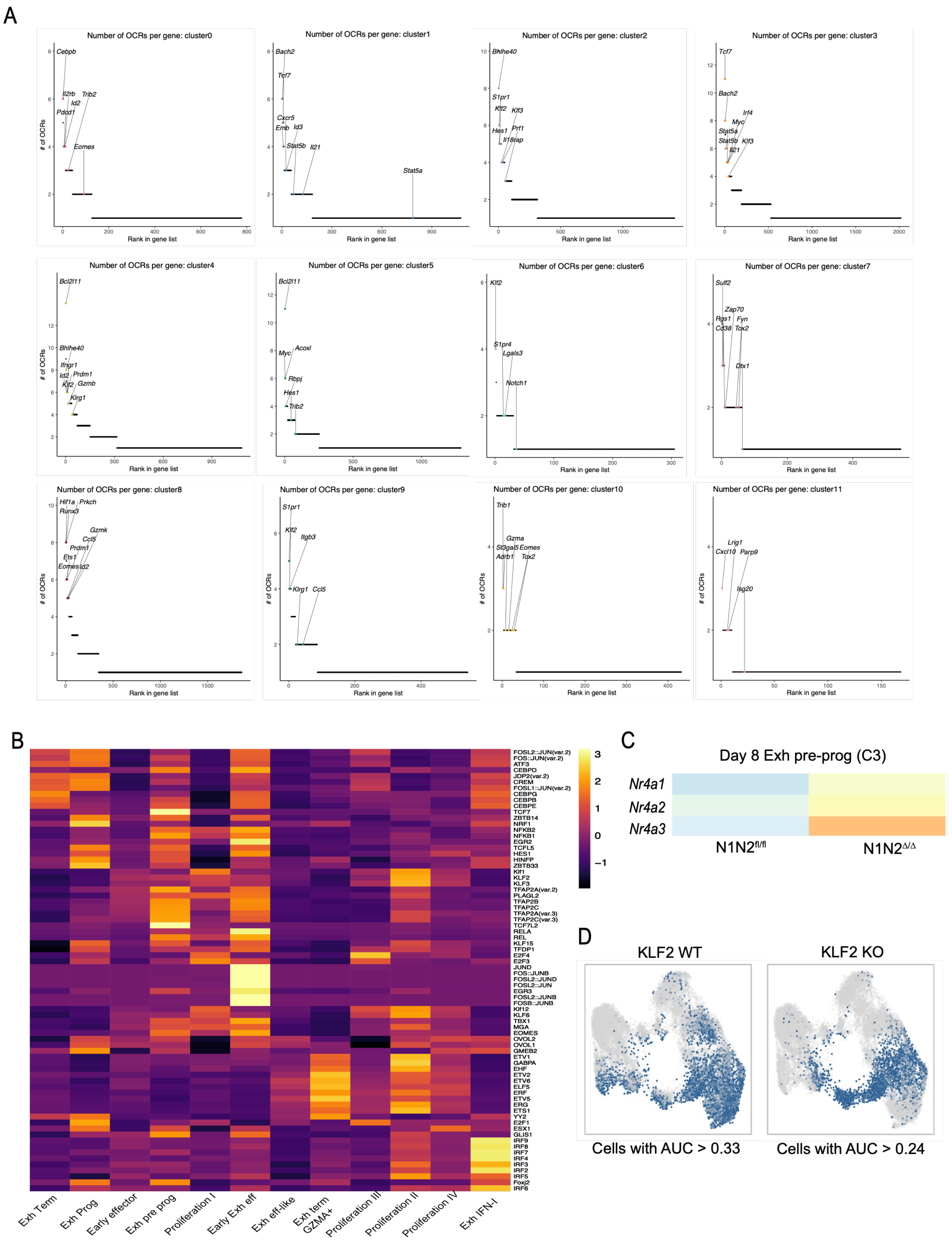
A
Number of OCRs per gene: cluster0
Number of OCRs per gene: cluster1
Number of OCRs per gene: cluster2
Number of OCRs per gene: cluster3
Number of OCRs per gene: cluster4
Number of OCRs per gene: cluster5
Number of OCRs per gene: cluster6
Number of OCRs per gene: cluster7
Number of OCRs per gene: cluster8
Number of OCRs per gene: cluster9
Number of OCRs per gene: cluster10
Number of OCRs per gene: cluster11
### of OCRs
Rank in gene list
B
Exh Term
Exh Prog
Early effector
Exh pre prog
Proliferation I
Early Exh eff
Exh eff-like
Exh term GZMA+
Proliferation III
Proliferation II
Proliferation IV
Exh IFN-I
C
Day 8 Exh pre-prog (C3)
Nr4a1
Nr4a2
Nr4a3
N1N2fl/fl
N1N2Δ/Δ
D
KLF2 WT
KLF2 KO
Cells with AUC > 0.33
Cells with AUC > 0.24

**Figure S5. Chromatin accessibility features associated with exhausted CD8$^+$ T cell clusters at day 8 and day 21 post–LCMV clone 13 infection.**

**(A)** Ranking of genes based on the number of associated open chromatin regions (OCRs) across individual exhausted CD8$^+$ T cell clusters identified by snRNA/ATAC-seq. For each cluster, genes are ordered according to their rank in the gene list (x-axis), and the corresponding number of OCRs per gene is shown (y-axis), highlighting cluster-specific regulatory programs and chromatin accessibility. **(B)** Heatmap depicting normalized motif accessibility scores across exhausted CD8$^+$ T cell clusters. Color scale indicates relative motif accessibility (z-score), revealing distinct epigenetic landscapes for each cluster. **(C)** Transcription of the TCR-induced genes, *Nr4a1-3*, by Exh pre-porg (cluster 3) cells from N1N2$^{fl/fl}$ and N1N2$^{\Delta/\Delta}$ mice at day 8 post-infection. **(D)** Enrichment of the KLF2 signature (Fargerberg et al. 2025) in Exh pre pro cells (cluster 3) and effector cells (clusters 2, 5 and 6).

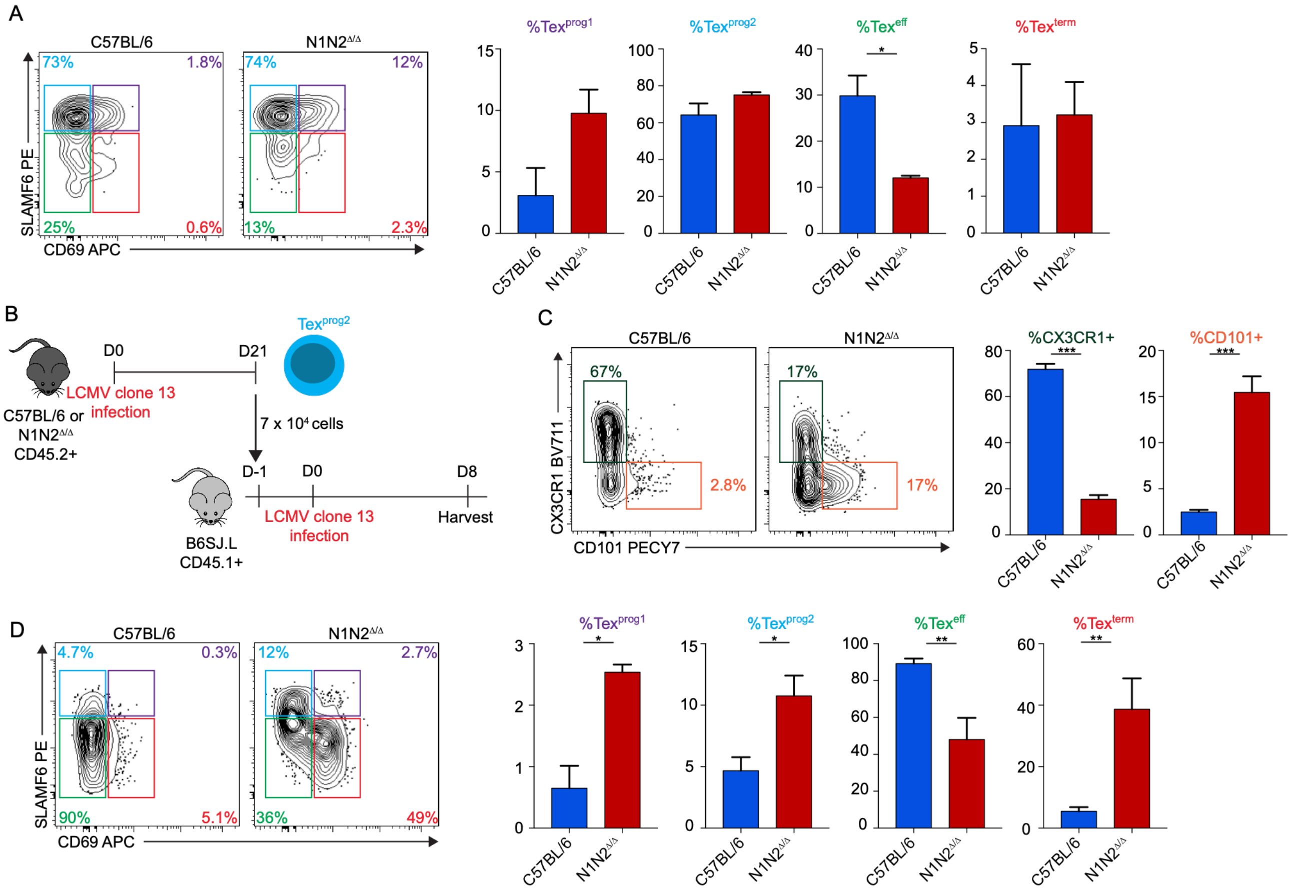


**Figure S6. Notch signaling regulates the differentiation of Tex progenitors into Tex effector-like cells after rechallenging with LCMV clone 13.**

**(A)** Adoptively transferred C57BL/6 and N1N2$^{\Delta/\Delta}$ Tex prog2 cells at day 21 post-infection into LCMV clone 13 match-infected B6.SJL recipient mice (CD45.1) (Fig. 6C) where we measured the expression of SLAMF6 and CD69 at day 7 post-transfer. (**B**) Experimental design for the adoptive transfer of day 21 Tex prog2 cells C57BL/6 or N1N2$^{\Delta/\Delta}$ into naive recipients (CD45.1+). Naïve mice were adoptively transferred with sorted cells and infected the next day with LCMV clone 13. Mice were euthanized 8 days post-infection. Proportion of CX3CR1+ cells or CD101+ cells (**C**) and of Tex subsets (**D**) within the adoptively transferred CD8$^{+}$CD45.2$^{+}$ cells from C57BL/6 and N1N2$^{\Delta/\Delta}$ cells.

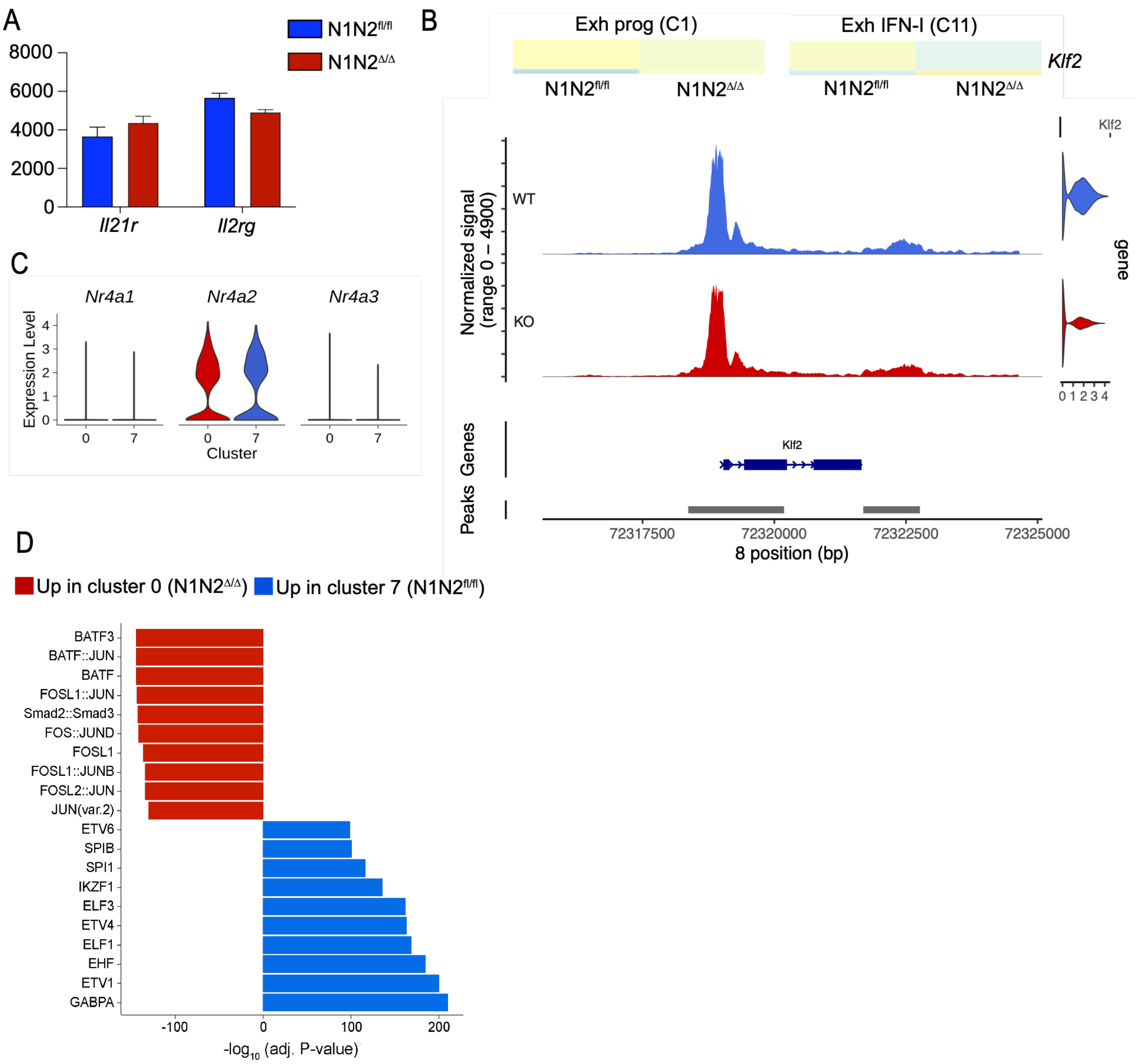


**Figure S7. The loss of Notch signaling during the differentiation of exhausted CD8⁺ T cells decreases *Klf2* transcription and gene accessibility while increasing TCR and TGF-β responses.**

**(A)** Notch signaling does not affect transcription of the genes coding for the IL-21 receptor. **(B)** Notch signaling regulates the transcription of *KLf2* and gene accessibility. **(C)** Notch-deficient Tex term cells transcribed more the TCR induced genes *Nr4a1*, *Nr4a2* and *Nr4a3*. **(D)** Smad2::Samd3 motif are enriched within the DARs of N1N2$^{\Delta/\Delta}$ Tex term cells (cluster 0) compared to their WT counterpart.
